# Additive energetic contributions of multiple peptide positions determine the relative promiscuity of viral and human sequences for PDZ domain targets

**DOI:** 10.1101/2022.12.31.522388

**Authors:** Elise F. Tahti, Jadon M. Blount, Sophie N. Jackson, Melody Gao, Nicholas P. Gill, Sarah N. Smith, Nick J. Pederson, Simone N. Rumph, Sarah A. Struyvenberg, Iain G. P. Mackley, Dean R. Madden, Jeanine F. Amacher

## Abstract

Protein-protein interactions that include recognition of short sequences of amino acids, or peptides, are critical in cellular processes. Protein-peptide interaction surface areas are relatively small and shallow, and there are often overlapping specificities in families of peptide-binding domains. Therefore, dissecting selectivity determinants can be challenging. PDZ domains are an example of a peptide-binding domain located in several intracellular signaling and trafficking pathways, which form interactions critical for the regulation of receptor endocytic trafficking, tight junction formation, organization of supramolecular complexes in neurons, and other biological systems. These domains are also directly targeted by pathogens, and a hallmark of many oncogenic viral proteins is a PDZ-binding motif. However, amidst sequences that target PDZ domains, there is a wide spectrum in relative promiscuity. For example, the viral HPV16 E6 oncoprotein recognizes over double the number of PDZ domain-containing proteins as the cystic fibrosis transmembrane conductance regulator (CFTR) in the cell, despite similar PDZ targeting-sequences and identical motif residues. Here, we determine binding affinities for PDZ domains known to bind either HPV16 E6 alone or both CFTR and HPV16 E6, using peptides matching WT and hybrid sequences. We also use energy minimization to model PDZ-peptide complexes and use sequence analyses to investigate this difference. We find that while the majority of single mutations had a marginal effect on overall affinity, the additive effect on the free energy of binding accurately describes the selectivity observed. Taken together, our results describe how complex and differing PDZ interactomes can be programmed in the cell.

## Introduction

Intracellular signaling and trafficking pathways are exquisitely tuned processes that depend on precise regulation and occur in a complex milieu. Both stable and dynamic protein-protein interactions are at the core of these biological systems. Of these, several, if not most, of the pathways include the recognition of conserved protein domains with short amino-acid sequences, also referred to as short linear motifs or peptides, on binding partners.^1–3^ These protein-peptide interactions tend to be of relatively low binding affinity and also often short-lived, allowing for a responsive environment where signals can be received and transmitted quickly.^4^ One such peptide-binding domain is the PSD-95/Dlg/ZO1 (PDZ) domain, named for the first three proteins identified as containing the conserved carboxylate-binding loop sequence that recognizes the extreme C-terminus of target proteins.^5–7^

PDZ domains are approximately 80-90 residues in length, with a conserved structural fold consisting of two *α*-helices and five or six *β*-strands, which form a core anti-parallel *β*-sheet.^8–11^ There are 272 PDZ domains in the human proteome within 154 proteins.^11^ These domains are implicated in the trafficking of membrane-bound receptors, organization of supramolecular structures, e.g., in the post-synaptic density of neurons, formation of tight junctions, and other critical processes.^12–15^ PDZ domains are also a target of pathogenic viral proteins, including examples from human papillomavirus (HPV), human immunodeficiency virus (HIV), influenza A, severe acute respiratory syndrome coronavirus 2 (SARS-CoV-2), and others.^16–27^ Specifically, analyses of several HPV strains indicate that those which contain PDZ domain recognition sequences in their E6 oncoproteins are cancer-causing, whereas strains lacking a PDZ-targeting E6 are not.^28^

Among these, the E6 oncoprotein from the HPV16 strain is particularly promiscuous for PDZ domains. It is known to bind over a dozen PDZ domain-containing proteins.^28^ Indeed, Vincentelli *et al*. found that *in vitro*, the final ten residues of the HPV16 E6 oncoprotein (sequence: SSRTRRETQL) bound 20% of the PDZome with *K*_D_ values < 250 μM using their holdup assay, which quantifies PDZ-peptide pairs for over 80% of known PDZ domains.^29, 30^ This is in contrast to known human sequences that target PDZ domains, e.g., the cystic fibrosis transmembrane conductance regulator (CFTR), which is recognized by relatively few.^31–34^

The endocytic recycling of CFTR is mediated by PDZ domain interactions, specifically by the Na^+^/H^+^ exchanger regulatory factor (NHERF) proteins and CFTR associated ligand (CAL).^33–35^ There may also be additional PDZ domains that interact with CFTR, e.g., those of mast2 microtubule associated serine/threonine kinase (MAST205) and sorting nexin 27 (SNX27).^31, 32^ Overall, the CFTR sequence (TEEEVQDTRL) is seemingly much less promiscuous than that of HPV16 E6, despite sharing the same Class I PDZ motif residues of Leu at the P^0^ (or C-terminal) position and Thr at P^-2^ (for CFTR, R=P^-1^, D=P^-3^, Q=P^-4^, etc.). Class I PDZ domains recognize the binding motif sequence X-S/T-X-*ϕ*_COOH_ (where *ϕ*=hydrophobic amino acids, typically I/L/V/F, and X=any amino acid), and overall, the majority of PDZ domain selectivity is known to occur at the final six (P^0^-P^-5^) positions, which were the focus of this work.^11, 36, 37^

Experiments investigating specificity in PDZ domains by ourselves and others previously found that target recognition is quite complex and dependent on position-specific interactions along the peptide-binding cleft.^36, 37^ Therefore, we were interested in investigating the basis of differences in PDZ recognition for the HPV16 E6 versus CFTR C-terminal sequences. We focused on the P^-1^, P^-4^, and P^-5^ positions, reasoning that the P^-3^ Glu and Asp, respectively, were chemically similar. Here, we describe our work determining binding affinities for nine PDZ domains that are known to bind HPV16 E6 alone, or CFTR and HPV16 E6 proteins, with peptides matching the WT sequences and variant peptides where we mutate one peptide position individually to the other sequence, e.g., the P^-1^, P^-4^, or P^-5^ CFTR residues into HPV16 E6 and *vice versa*. We use structural models and sequence analyses to investigate clear amino-acid preferences where the mutation changes the |*ΔΔ*G°| of binding by more than 1 kcal/mol in either direction, as well as determine that in general, polar residues are preferred at these surface-exposed sites by a majority of PDZ domains.

We conclude that while the P^-1^ Arg and P^-5^ Val amino acids in CFTR most strongly contribute to its relatively restricted PDZ selectivity, a single mutation only marginally affects binding affinity in the majority of the protein-peptide pairs studied. Instead, it is the additive energetic contributions from all three positions that result in a sequence which binds below a threshold that is endogenously relevant. Our work is likely applicable broadly across the PDZ family, providing insight into how the overlapping yet distinct PDZ interactomes are encoded in domains of these important cellular pathways.

## Results

### I. The effect of single position variants on HPV16 E6 binding for NHERF1 PDZ1 (N1P1), NHERF2 PDZ2 (N2P2), CAL PDZ, or TIP-1 PDZ domains

To investigate the position-specific characteristics of PDZ domain binding to sequences matching the C-termini of the CFTR and HPV16 E6 proteins, we initially chose a subset of PDZ domains that bind HPV16 E6 and either target CFTR with high affinity (N1P1 and N2P2), low affinity (CAL), or not at all (TIP-1).^18, 38–40^ We previously studied this subset of PDZ domains with respect to their ability to bind CFTR and/or a designed inhibitor of CAL, iCAL36.^41^ The NHERF PDZ domains, including N1P1 and N2P2, bind a peptide matching the CFTR C-terminal sequence (sequence: TEEEVQDTRL) with sub-micromolar affinity, whereas previous fluorescence polarization assays showed CAL binds this sequence with *K*_i_ = 420 ± 80 μM and TIP-1 shows undetectable binding, defined as >1000 μM.^41^ Notably, CAL also binds an HPV16 E6 sequence (SSRTRRETQL) with relatively low affinity, *K*_i_ = 340 ± 70 μM.^36^

We designed a set of five peptides to test which, if any, amino acids in the CFTR sequence contribute to these relative binding affinities. These included: HPV16 E6 (defined above), _SSRT_CFTR (SSRTVQDTRL), HPV_P1R_ (SSRTRRETRL), HPV_P4Q_ (SSRTRQETQL), and HPV_P5V_ (SSRTVRETQL). In each of the HPV16 E6 variant peptides, the indicated position, P^-1^, P^-4^, or P^-5^, respectively, is mutated to the corresponding amino acid in the CFTR sequence. We chose not to include the P^—3^ variant peptide, as we reasoned that Glu (in HPV16 E6) and Asp (in CFTR) are chemically very similar. We included the P^-9^-P^-6^ SSRT amino acids in all sequences, including CFTR, for consistency. Decameric peptides were used due to previous work showing a modest affinity increase with sequences as long as 10 residues and N1P1.^34^

Binding affinities for this set of PDZ domains and peptides were determined using fluorescence polarization competition experiments. All proteins were expressed and purified, and assays conducted, as described previously and in the Materials and Methods.^34, 36, 41^ The binding affinities and free energies determined from triplicate experiments are in **Table 1**.

**Table 1.** Binding affinity data for N1P1, N2P2, CAL, and TIP-1 bound to HPV16 E6, _SSRT_CFTR, and HPV variant peptides (HPV_P1R_, HPV_P4Q_, and HPV_P5V_).

| | $K_i$ (μM) (in parentheses, $\Delta G^\circ$ in kcal/mol) | | | | |
| --- | --- | --- | --- | --- | --- |
|  | HPV16 E6 | HPV<sub>P1R</sub> | HPV<sub>P4Q</sub> | HPV<sub>P5V</sub> | SSRTCFTTR |
| N1P1 | 100 ± 50<br>(-5.49) | 8.8 ± 3.4<br>(-6.94) | 21 ± 7<br>(-6.42) | 67 ± 28<br>(-5.73) | 2.2 ± 1<br>(-7.77) |
| N2P2 | 17 ± 6<br>(-6.55) | 6.9 ± 1.1<br>(-7.08) | 5.2 ± 0.6<br>(-7.25) | 7.0 ± 3.6<br>(-7.08) | 1.7 ± 0.4<br>(-7.92) |
| CAL | 160 ± 20<br>(-5.18) | 42 ± 10<br>(-5.97) | 56 ± 6<br>(-5.80) | 190 ± 10<br>(-5.07) | 130 ± 10<br>(-5.30) |
| TIP-1 | 11 ± 1<br>(-6.76) | 22 ± 5<br>(-6.35) | 34 ± 2<br>(-6.09) | 35 ± 4<br>(-6.08) | 670 ± 400<br>(-4.33) |

Despite having similar affinities for _SSRT_CFTR (1.7 ± 0.4 μM for N2P2, 2.2 ± 1 μM for N1P1), the binding affinity of N2P2 was significantly higher than that of N1P1 for the HPV16 E6 peptide: 17 ± 6 μM versus 100 ± 50 μM, respectively. For N1P1 and N2P2, mutating the P^-1^, P^-3^, or P^-5^ residues in the HPV16 E6 sequence to the corresponding amino acids in CFTR increased the relative binding affinity (**Table 1, Figures 1A-B**). In N1P1, the P^-1^ Arg in HPV_P1R_ increased the relative binding affinity the greatest, at >10-fold, whereas the effect was more modest for a P^-4^ Gln (HPV_P4Q_) at 5-fold, and minimal for a P^-5^ Val (HPV_P5V_) (**Table 1**). For N2P2, all 3 mutations in HPV16 E6 resulted in a similar increase in affinity, of 2.4-3.3-fold (**Table 1, Figure 1B**). The preference of NHERF PDZ domains for a P^-1^ Arg is known, and attributed to an observed electrostatic interaction between P^-1^ Arg on the peptide and Glu43 on the protein.^42, 43^ In N2P2, the equivalent P^-1^ Arg-interacting E43 residue is D180.

**Figure 1.**
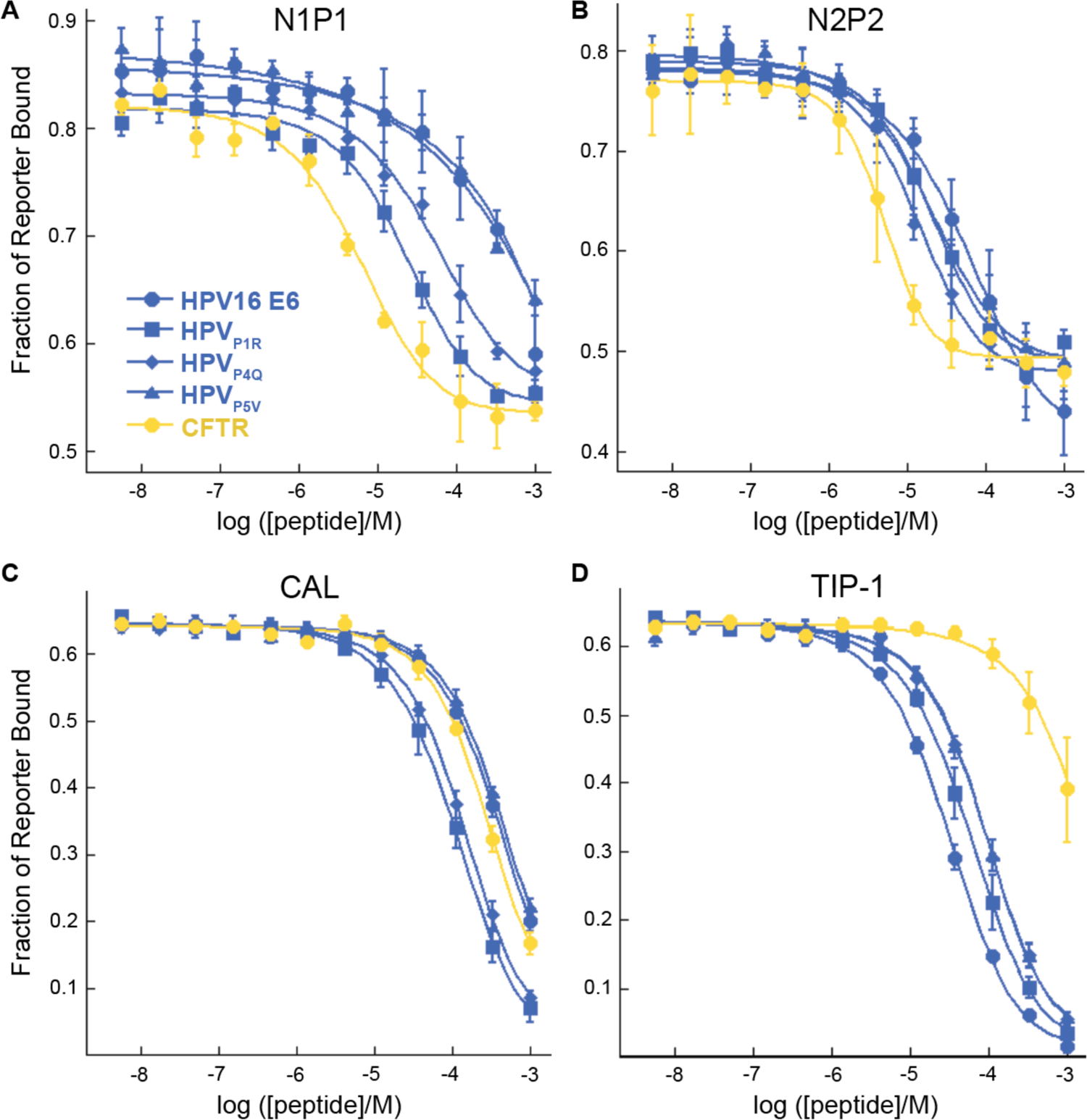
Competition binding curves for N1P1, N2P2, CAL, and TIP-1 with _SSRT_CFTR, HPV16 E6, and HPV variant peptides (HPV_P1R_, HPV_P4Q_, HPV_P5V_). The average results of at least triplicate experiments are shown, with error bars reflecting standard deviation. Reporter peptides used are described in the Materials and Methods. Curves were visualized using Kaleidagraph, using previously reported methods (see main text).

In CAL, which binds both HPV16 E6 and CFTR relatively weakly, at >100 μM, there was no change in overall binding affinity for a P^-5^ Val mutation, with *K*_i_ = 190 ± 10 μM (**Table 1, Figure 1C**). However, mutating either the P^-1^ HPV16 E6 Gln residue to Arg or the P^-4^ Arg to Gln increased the relative affinity of CAL, as compared to HPV16 E6, by 3.8- and 2.9-fold, respectively (**Table 1, Figure 1C**). Previous peptide-array data, with single-position mutations to all other amino acids for 10 peptide sequences, revealed that a P^-1^ Arg is a positive modulator for CAL. At P^-4^, both a Gln and Arg are favorable and equivalently tolerated within these peptide contexts.^36^ Therefore, it is interesting that the HPV_P4Q_ peptide revealed an almost 3-fold higher affinity for CAL (**Table 1**). Although not studied here, one idea based on the 4K75 structure, where the P^-4^ Gln interacts with E308 in the βB-βC loop, is that the bulky P^-4^ Arg is unable to make this same interaction. This result may also suggest that sequence context can affect modulator (or non-motif) preferences for PDZ domains under certain conditions.

For TIP-1, binding to CFTR was weakly detectable, with *K*_i_ = 670 ± 400 μM for the _SSRT_CFTR sequence, slightly better than the previously calculated value for WT CFTR, likely due to the SSRT N-terminal substitution.^41^ Furthermore, no single point mutation in HPV16 E6 to the corresponding amino acid in CFTR dramatically affected the overall binding affinity (**Table 1, Figure 1D**). While each individual mutation revealed an approximately 2-3-fold effect weakening of binding affinity relative to the WT HPV16 sequence, the combination resulted in 60-fold lower binding for _SSRT_CFTR, as compared to HPV16 E6. The _SSRT_CFTR sequence also contains the P^-3^ Glu to Asp substitution, which was not tested here.

Overall, the largest effect seen in these PDZ-peptide pairs was for N1P1 and HPV_P1R_, which increased the binding affinity by >10-fold as compared to the HPV16 E6 sequence. The *ΔΔ*G° value for this mutation = −1.45 kcal/mol (**Figure 2**). All other affinity changes were relatively modest, with |*ΔΔ*G°| values, as compared to HPV16 E6, less than 1 kcal/mol. Taken together, this suggested that single point mutants do not make a substantial contribution to the overall free energy of binding (**Figure 2**).

**Figure 2.**
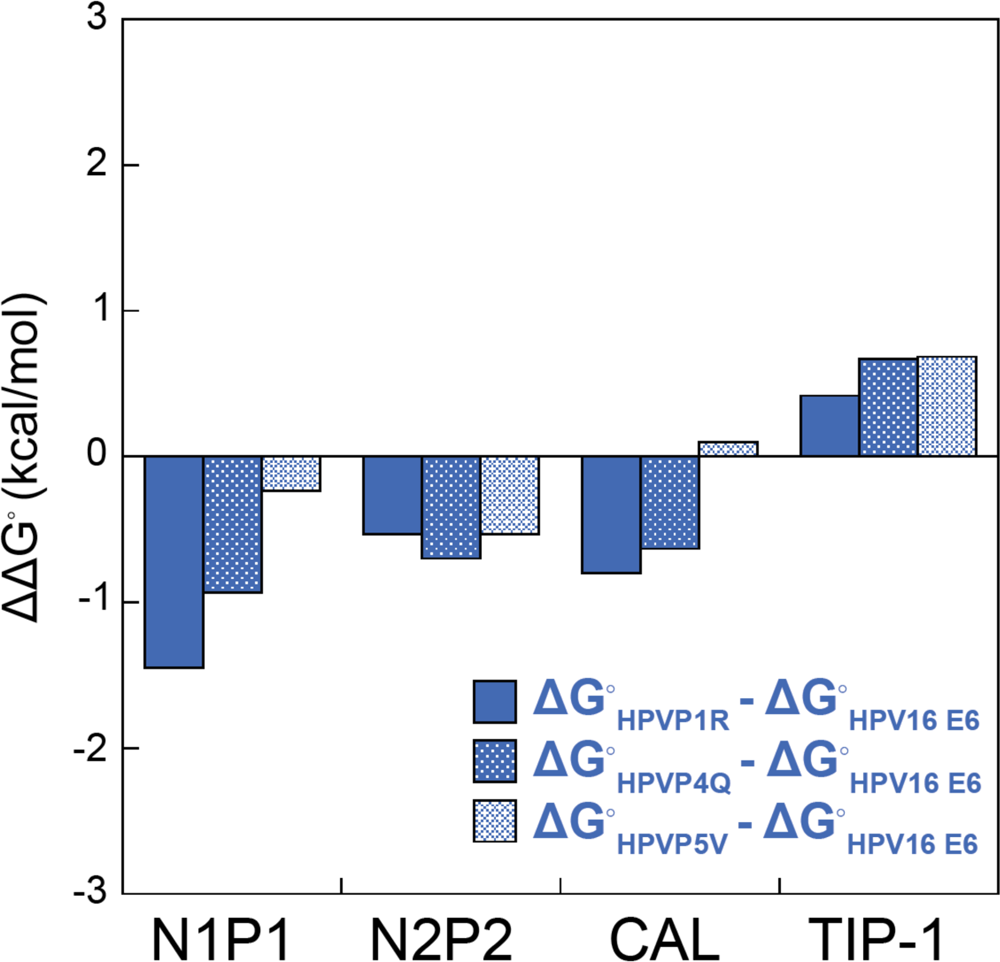
ΔΔG° values (in kcal/mol) for N1P1, N2P2, CAL, and TIP-1 with _SSRT_CFTR, HPV16 E6, and HPV variant peptides (HPV_P1R_, HPV_P4Q_, HPV_P5V_). ΔG° values were calculated using the equation *R*T*ln(*K*_i_), where *R*=0.001987 kcal mol^-1^ K^-1^, T=300 K, and the *K*_i_ values are in **Table 1**.

### II. Binding affinities of PDZ domains that target the HPV16 E6 sequence are relatively resistant to single point variants in either the HPV16 E6 or CFTR sequences

Our TIP-1 results were of particular interest, showing relatively minor changes, *≤* 3-fold, in overall binding affinity upon mutation of the HPV16 E6 P^-1^, P^-4^, or P^-5^ positions, despite a 60-fold difference in affinity of the WT sequence versus CFTR. We next wanted to test if other HPV16 E6-targeting PDZ domains would show a similar result, whereby site-specific substitutions of amino acids in the CFTR sequence would fail to dramatically decrease relative binding affinity. We chose several representative PDZ domains known to bind the HPV16 E6 oncoprotein, including: DLG1 PDZ1 (DLG1-1), MAGI1 PDZ2 (MAGI1-2), PTPN3 PDZ (PTPN3), SCRIB PDZ1 (SCRIB-1), and SNX27 PDZ (SNX27).^23, 27, 28, 44–49^ There are available structures of MAGI1-2 and PTPN3 bound to the HPV16 E6 sequence.^46, 50^

We recombinantly expressed and purified all PDZ domains as described in the Materials and Methods, and followed a similar protocol used with the PDZ domains described above. In addition to using fluorescence polarization to determine relative binding affinities for the HPV16 E6, _SSRT_CFTR, and HPV16 E6 variant peptides (HPV_P1R_, HPV_P4Q_, and HPV_P5V_) above, we also created analogous _SSRT_CFTR variant peptides, with substitutions at the P^-1^, P^-4^, and P^-5^ positions to amino acids in the HPV16 E6 sequence. These peptides are: CFTR_P1Q_ (SSRTVQDTQL), CFTR_P4R_ (SSRTVRDTRL), and CFTR_P5R_ (SSRTRQDTRL). We tested the CFTR variant peptides with N1P1 and N2P2 as well.

In order to conduct competition experiments using fluorescence polarization, we first determined *K*_D_ values for a fluoresceinated HPV16 E6 reporter peptide (*FITC*-SSRTRRETQL) with DLG1-1, MAGI1-2, PTPN3, and SCRIB-1. We previously determined that SNX27 binds F*-GIRK3 (*FITC*-LPPPESESKV) with *K*_D_ = 0.33 ± 0.14 μM.^51^ Calculated *K*_D_ values for triplicate experiments were: DLG1-1 (17 ± 4 μM), MAGI1-2 (12 ± 4 μM), PTPN3 (2 ± 1 μM), and SCRIB-1 (5.1 ± 1.7 μM) (**Figure 3A-D**).

**Figure 3.**
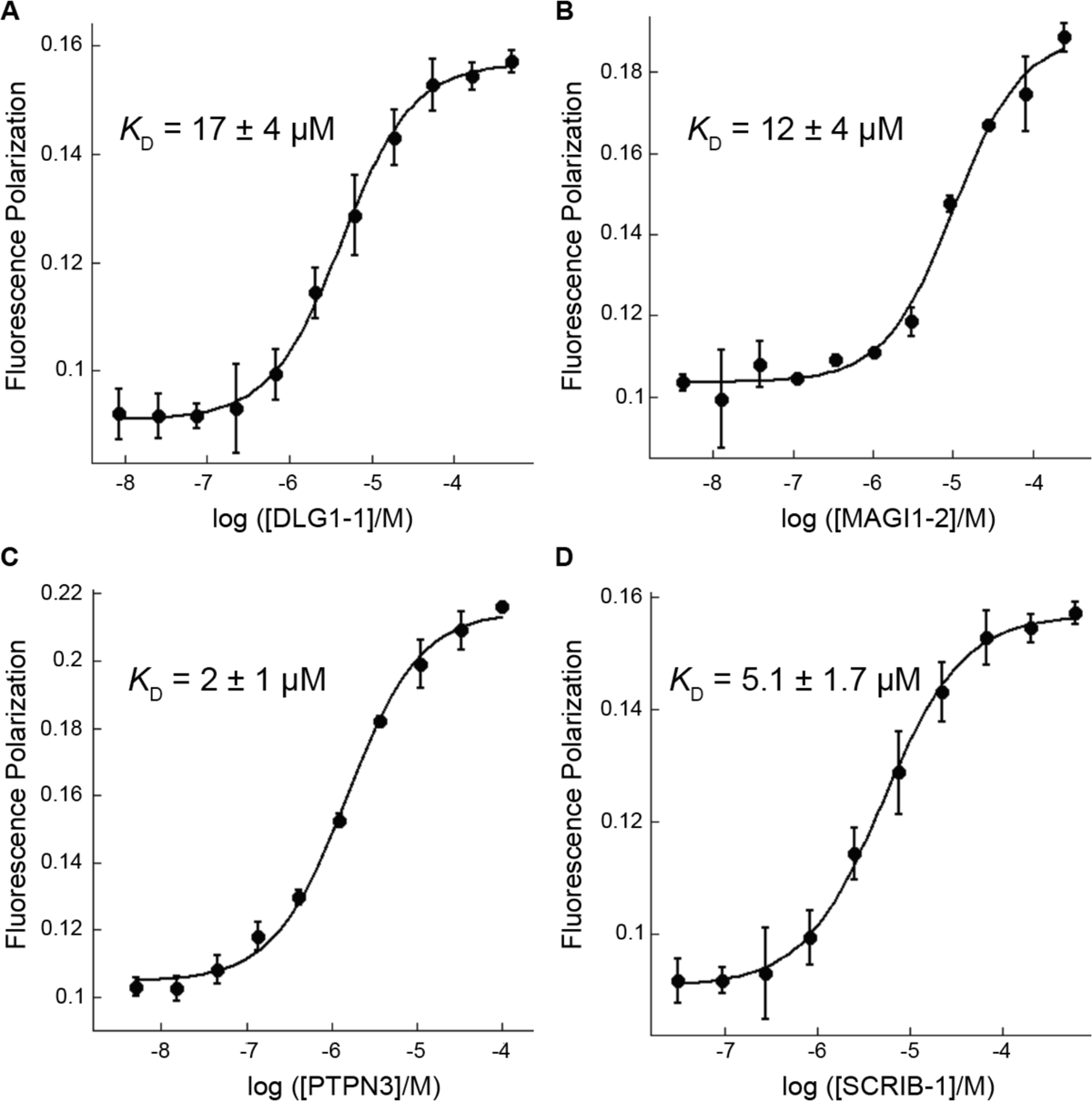
*K*_D_ curves for PDZ domain-*F*\*HPV16 E6 reporter peptide experiments. Although *K*_D_ values were determined from triplicate experiments, here the average and standard deviation for N=3 is shown due to a difference in protein concentration range. *K*_D_ values were determined using Kaleidagraph, using previously reported methods (see main text).

Competition experiments with our set of eight peptides (HPV16 E6, _SSRT_CFTR, and the P^-1^, P^-4^, P^-5^ HPV16 E6 and CFTR variants) generally agreed with our previous results, with a few exceptions (**Table 2, Figure 4**). Again, we calculated the free energy of binding for each PDZ-peptide pair, as well as *ΔΔ*G° values as compared to HPV16 E6 or CFTR (**Figure 5A**). Free energies for interactions > 1 mM were calculated using *K*_i_ = 1000 μM, and likely underestimate the true values. For MAGI1-2, the *K*_D_ value was too high compared to the *K*_i_ values and calculations of *K*_i_ using our approach were not successful. For PTPN3, calculations of *K*_i_ were also not successful, likely because we overestimated the amount of protein ideally used in competition experiments. In both cases, we calculated IC_50_ values, based on the concentration of inhibitor peptide at half-maximal saturation (**Table 2, Figure 4F-G**). The IC_50_ values were used to approximate *Δ*G° and *ΔΔ*G° for these PDZ domains.

**Figure 4.**
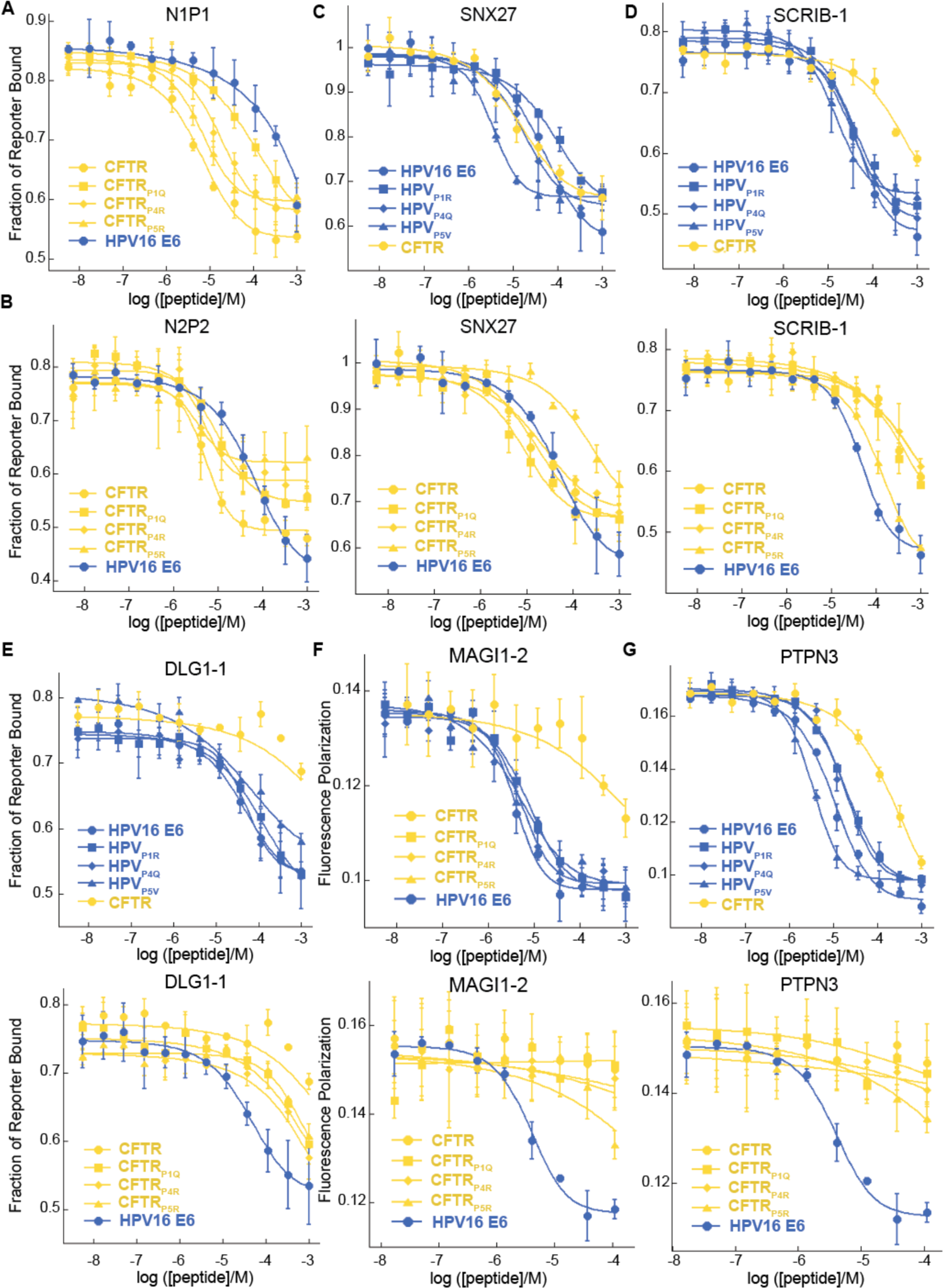
Competition binding experiments for PDZ domains studied with HPV16 E6, HPV variants (HPV_P1R_, HPV_P1Q_, HPV_P5V_), _SSRT_CFTR, and CFTR variants (CFTR_P1Q_, CFTR_P4R_, CFTR_P5R_). All experiments are at least triplicate experiments. For DLG1-1, only N=2 data are shown due to higher standard deviations. All data are included in **Table 2**. (**F-G**) These data are shown as a function of fluorescence polarization due to the inability to accurately calculate the fraction of reporter bound for the MAGI1-2 and PTPN3 PDZ domains.

**Figure 5.**
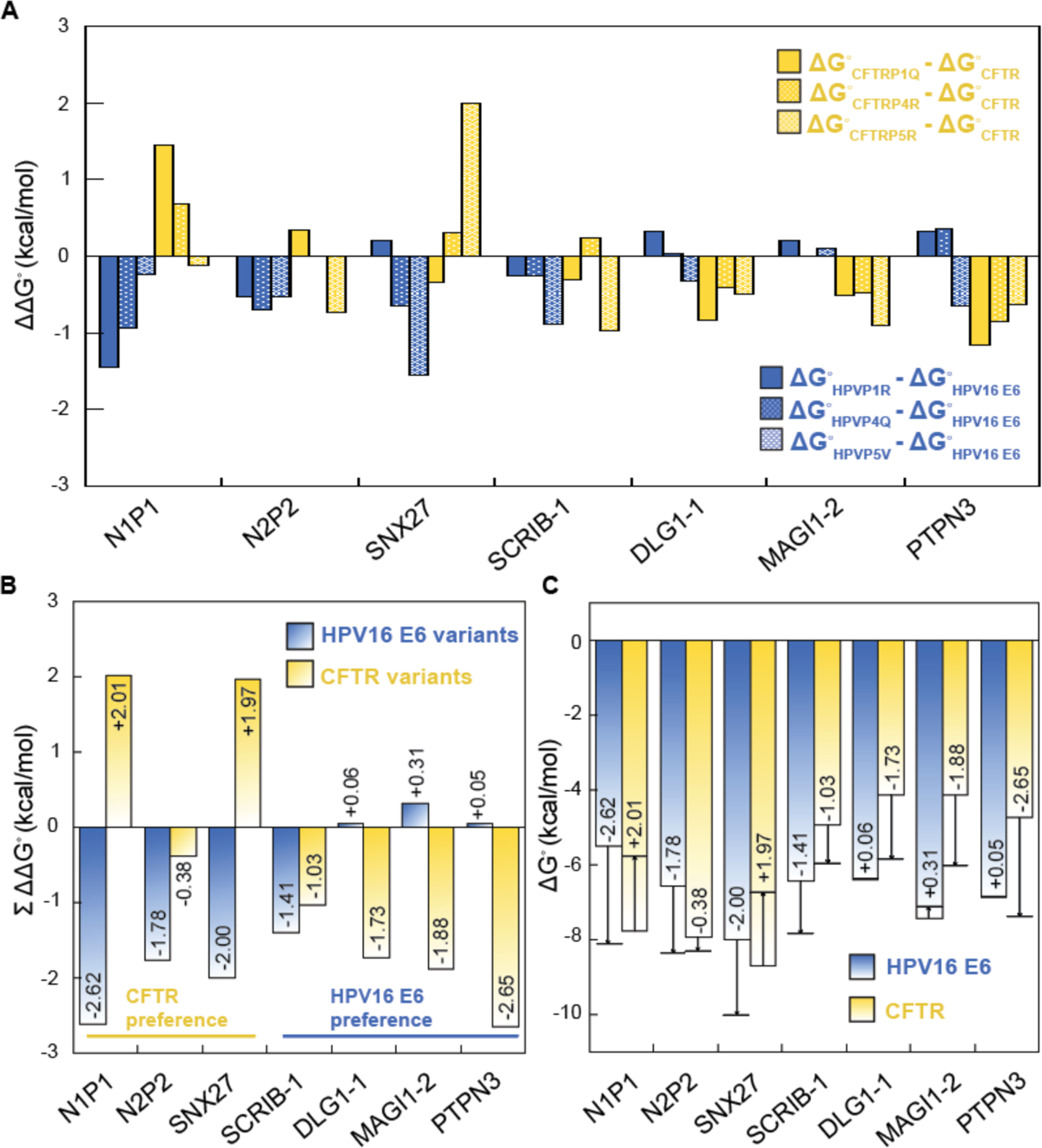
ΔΔG° and Σ ΔΔG° values (in kcal/mol) for binding experiments in Table 2. **(A)** ΔΔG° values calculated as in Figure 2. (**B**) *Σ ΔΔ*G° values were calculated as the sum of the HPV variant or CFTR variant ΔΔG° values in (**A**). Here, a higher -Σ ΔΔG° is more favorable, so e.g., N1P1, N2P2, and SNX27 show a preference for the CFTR sequence, considering mutations in HPV16 E6 to CFTR residues result in a large -ΣΔΔG°. DLG1-1, MAGI1-2, and PTPN3 strongly prefer HPV16 E6. The data for SCRIB-1 is less clear, but **Table 2** confirms that this PDZ domain does prefer HPV16 E6, as this representation fails to capture quantitative magnitude of binding affinities. (**C**) The black arrows indicate Σ ΔΔG° values (from **B**) added to the ΔG° of binding for each PDZ domain to either HPV16 E6 (blue) or CFTR (yellow).

**Table 2.** Binding affinity data for N1P1, N2P2, DLG1-1, SCRIB-1, SNX27, MAGI1-2, and PTPN3 bound to HPV16 E6, _SSRT_CFTR, HPV variant peptides (HPV_P1R_, HPV_P4Q_, and HPV_P5V_), and CFTR variant peptides (CFTR_P1Q_, CFTR_P4R_, CFTR_P5R_). All data are the result of at least triplicate experiments. The free energies of binding are in parenthesis (in kcal/mol).

| | $K_i$ (μM) (in parentheses, $\Delta G^\circ$ in kcal/mol) | | | | | | | |
| --- | --- | --- | --- | --- | --- | --- | --- | --- |
|  | HPV16 E6 | HPV<sub>P1R</sub> | HPV<sub>P4Q</sub> | HPV<sub>P5V</sub> | SSRTCFTR | CFTR<sub>P1Q</sub> | CFTR<sub>P4R</sub> | CFTR<sub>P5R</sub> |
| N1P1<sup>a</sup> | 100 ± 50<br>(-5.49) | 8.8 ± 3.4<br>(-6.94) | 21 ± 7<br>(-6.42) | 67 ± 28<br>(-5.73) | 2.2 ± 1<br>(-7.77) | 25 ± 7<br>(-6.32) | 6.9 ± 2.7<br>(-7.08) | 1.8 ± 0.3<br>(-7.89) |
| N2P2<sup>a</sup> | 17 ± 6<br>(-6.55) | 6.9 ± 1.1<br>(-7.08) | 5.2 ± 0.6<br>(-7.25) | 7.0 ± 3.6<br>(-7.08) | 1.7 ± 0.4<br>(-7.92) | 3.1 ± 0.5<br>(-7.56) | 1.7 ± 0.5<br>(-7.92) | 0.49 ± 0.38<br>(-8.66) |
| DLG1-1 | 22 ± 12<br>(-6.39) | 39 ± 11<br>(-6.05) | 23 ± 6<br>(-6.37) | 13 ± 4<br>(-6.71) | >1000<br>(-4.12) | 250 ± 150<br>(-4.94) | 510 ± 230<br>(-4.52) | 430 ± 200<br>(-4.62) |
| SCRIB-1 | 20 ± 7<br>(-6.45) | 13 ± 2<br>(-6.71) | 13 ± 2<br>(-6.71) | 4.5 ± 2.8<br>(-7.34) | 250 ± 30<br>(-4.94) | 150 ± 60<br>(-5.25) | 370 ± 170<br>(-4.71) | 50 ± 10<br>(-5.90) |
| SNX27 | 1.5 ± 0.5<br>(-7.99) | 2.1 ± 0.7<br>(-7.79) | 0.51 ± 0.22<br>(-8.64) | 0.11 ± 0.03<br>(-9.55) | 0.46 ± 0.11<br>(-8.70) | 0.26 ± 0.05<br>(-9.04) | 0.44 ± 0.06<br>(-8.72) | 7.4 ± 4.5<br>(-7.04) |
| | $IC_{50}$ (μM) (in parentheses, $\Delta G^\circ$ in kcal/mol) | | | | | | | |
| MAGI1-2 | 3.7 ± 0.1<br>(-7.45) | 5.2 ± 2.4<br>(-7.25) | 3.8 ± 2.1<br>(-7.44) | 4.4 ± 1.1<br>(-7.35) | >1000<br>(-4.12) | 420 ± 210<br>(-4.63) | 460 ± 200<br>(-4.59) | 220 ± 50<br>(-5.02) |
| PTPN3 | 10 ± 1<br>(-6.86) | 18 ± 1<br>(-6.52) | 18 ± 5<br>(-6.50) | 3.4 ± 0.3<br>(-7.51) | 350 ± 250<br>(-4.74) | 51 ± 14<br>(-5.89) | 85 ± 39<br>(-5.59) | 120 ± 20<br>(-5.37) |

<sup>a</sup>N1P1 and N2P2 values with HPV16 E6 and HPV variant peptides are also reported in **Table 1**.

Surprisingly, SNX27 PDZ bound CFTR with high affinity, better than its affinity for HPV16 E6 and all variants, excluding HPV16_P5V_ (**Table 2, Figure 4C**). SNX27 was previously implicated in *β*_2_-adrenoreceptor (*β*_2_AR) PDZ-mediated recycling from early endosomes, and may play a similar role in CFTR trafficking.^52^ This hypothesis is directly supported by a pre-print article that determined SNX27 does indeed directly bind and mediate the trafficking of CFTR.^32^

Overall, there were seven PDZ-peptide interactions that revealed |*ΔΔ*G°| values > 1 kcal/mol. These included: N1P1 (HPV_P1R_ - HPV16 E6 and CFTR_P1Q_ - _SSRT_CFTR), SCRIB-1 (HPV16_P4Q_ and HPV16_P5V_ - HPV16 E6), SNX27 (HPV_P5V_ - HPV16 E6 and CFTR_P5R_ - _SSRT_CFTR) and PTPN3 (CFTR_P1Q_ - _SSRT_CFTR) (**Tables 2-3, Figure 5A**). Notably, for the P^-1^ position with N1P1 and P^-5^ position with SNX27, the variant peptides showed effectively equal and opposite *ΔΔ*G° values: N1P1 (−1.45 kcal/mol for HPV_P1R_ - HPV16 E6 and +1.45 kcal/mol for CFTR_P1Q_ - _SSRT_CFTR) and SNX27 (−1.56 kcal/mol for HPV_P5V_ - HPV16 E6 and +2.0 kcal/mol for CFTR_P5R_ - _SSRT_CFTR). This suggests strong preferences for N1P1 and a P^-1^ Arg, as previously discussed, as well as SNX27 and a P^-5^ Val.

**Table 3.**
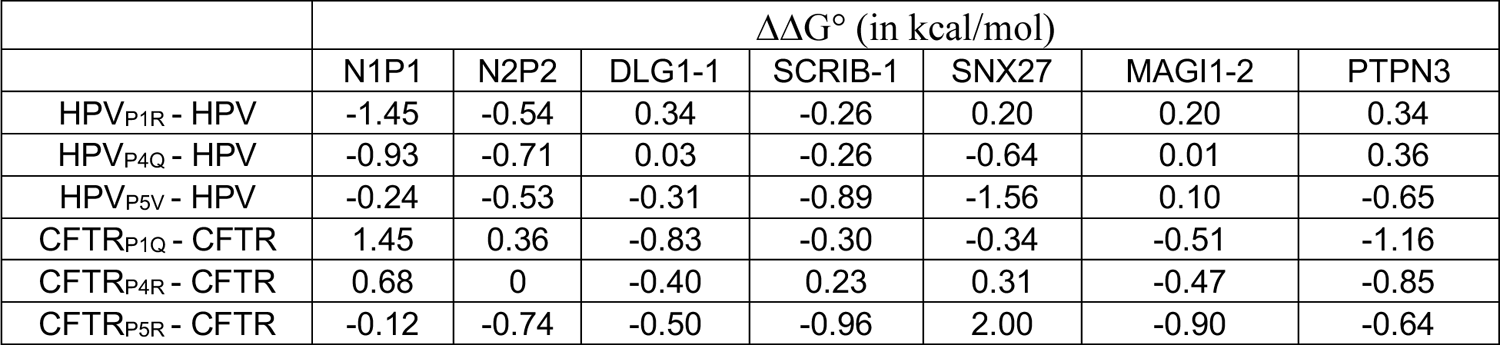
ΔΔG° values for PDZ-peptide pairs. ΔΔG° values were calculated as the difference between the variant peptide and WT peptide (HPV16 E6 or _SSRT_CFTR) sequence.

Overall, of 48 *ΔΔ*G° values calculated, only seven exceeded ± 1 kcal/mol, of which five have binding affinity changes >10-fold. Another 19 resulted in |*ΔΔ*G°| > 0.5 kcal/mol. DLG1-1, MAGI1-2, and PTPN3 were generally characteristic of TIP-1, whereby mutations of HPV16 E6 positions to amino acids in CFTR decrease binding affinity, and although there are exceptions (DLG1-1 and HPV_P5V_, PTPN3 and HPV_P5V_), they are relatively modest (**Table 2, Figure 5A**). In addition, for these 3 PDZ domains, substitution of CFTR positions to amino acids in HPV16 E6 increased relative binding affinity in all cases (**Table 2, Figure 5A**). Taken together, these peptide positions clearly modulate affinity and selectivity for the tested PDZ domains. However, in general, overall binding for a given sequence was relatively resistant to single amino-acid substitutions, even to those from a sequence that it binds poorly or not at all.

Finally, we calculated the *Σ ΔΔ*G° values for either the HPV16 E6 variant peptides, as compared to HPV16 E6, or the CFTR variant peptides, as compared to _SSRT_CFTR (**Figures 5B-C**). In most cases, this revealed the clear selectivity preference for the PDZ domain. For example, the additive effect for all three of the singly mutated CFTR variant peptides exceeded −1.5 kcal/mol for DLG1-1, MAGI1-2, and PTPN3, reflecting the clear preference of those PDZ domains for HPV16 E6. The additive effect for the HPV16 E6 variant peptides exceeded −1.5 kcal/mol for N1P1, N2P2, and SNX27, and indeed, our binding assays resulted in 3- to 45-fold higher affinity for _SSRT_CFTR than HPV16 E6 (**Table 2**).

The SCRIB-1 PDZ domain was an interesting case out of this test set; although there was a clear preference for HPV16 E6 over CFTR, with *K*_i_ = 20 μM and 250 μM, respectively, mutation of the P^-5^ position in the variant peptides showed a 5-fold enhancement of affinity in both background sequences. As a result, the additive effect of these reciprocal mutations in both HPV16 E6 and CFTR backbones was a favorable *Σ ΔΔ*G° that exceeded −1 kcal/mol. However, even the highest affinity CFTR variant for SCRIB-1, CFTR_P5R_ was a 2.5-fold worse binder than HPV16 E6, confirming the clear preference for HPV16 E6 (**Table 2, Figure 5C**). It would be interesting to test doubly mutated peptides with SCRIB-1 in future experiments to explore the context dependence of these mutations. In general, these additive free energies of binding are consistent with the affinity differences of each domain for HPV16 E6 and CFTR.

### III. Energy minimizations used to analyze position-specific substitutions in structural models

To understand the stereochemistry of our binding assay results, we analyzed structural models of these PDZ-peptide complexes. Three of the PDZ/peptide pairs studied have available structures, including N1P1/CFTR (PDB 1I92), MAGI1-2/HPV16 E6 (6TWQ), and PTPN3/HPV16 E6 (6HKS).^42, 46, 50^ Other structures used were: DLG1-1 (3RL7), N2P2 (2HE4), SCRIB-1 (6MYF), and SNX27 (3QGL).^53–56^

We used PyMOL to model P^0^-P^-5^ of the HPV16 E6 and CFTR sequences in the PDZ domains without experimental structures. The HPV16 E6 peptide from PDB 6TWQ (MAGI1-2) was used to create initial structural models of N1P1/HPV16 E6, N2P2/HPV16 E6, DLG1-1/HPV16 E6, and SCRIB-1/HPV16 E6 via direct alignment. For the CFTR peptide, the HPV16 E6 peptide was mutated using the PyMOL mutagenesis wizard for all PDZ domains. Analysis of the P^-4^ Gln and P^-5^ Val residues in the N1P1/CFTR complex structure suggests that the P^-4^ Gln binds in the traditional P^-5^ pocket on the PDZ domain, and consequently, the P^-5^ residue, a Glu in N1P1 (1I92), is interacting with solvent (**Figure S1A**). This is likely due to the constrained geometry of this crystal structure, as N1P1 binds a C-terminal extension in a molecule related by symmetry that matches the CFTR sequence.^42^ For our analyses, we chose to mutate the HPV16 E6 sequence instead of using this conformation. Finally, the SNX27/HPV16 E6 and SNX27/CFTR models were created by mutating GIRK3 (ESESKV) amino acids *in silico*, due to shifts of 2.7-4.9 Å between P^-5^ C*_α_* atoms and DLG1-2/HPV18 E6 (PDB 2OQS, used as an additional comparison here), PTPN3/HPV16 E6, and MAGI1-2/HPV16 E6, respectively (**Figure S1B**).

PDZ-peptide binding is well understood, with dozens of available structures.^11^ However, there were clear steric clashes and other minor issues with our initial models. Therefore, we ran energy minimization on a solvated system to refine our models for analysis, as described in the Materials and Methods. We reasoned that full molecular dynamics simulations would be unlikely to inform much new knowledge for these complexes, based on prior PDZ knowledge, and that this approach would provide a reasonable structural model analogous to docking simulations.

Alignment of the HPV-binding structures and models reveals position-specific recognition by PDZ residues. There are only two specific interactions between P^-1^ Gln and PDZ residues, including N1P1 H157 (3.1 Å between the side chain and imidazole ring) and SCRIB-1 R762 (3.6 Å between the side chain and guanidino group) (**Figure 6A**). In the other structures, there is a conserved polar residue at the *β*B-2 residue, immediately following the carboxylate binding loop motif, which is in the vicinity of the P^-1^ peptide position. In PTPN3, R539 is in the equivalent SCRIB-1 R762 position, however it is not directly interacting with the P^-1^ Gln. This is notable as the PTPN3/HPV16 E6 complex is an experimental structure, whereas our SCRIB-1/HPV16 E6 is a model. In PTPN3 PDZ, the side-chain amino group of K520 in the *β*A-*β*B loop is 5.4 Å from the side chain of P^-1^ Gln (**Figure 6A**). In general, all PDZ domains except SNX27 contain multiple hydrophilic residues in this region. SNX27 displays the most hydrophobic character, specifically due to A81 and L83.

**Figure 6.**
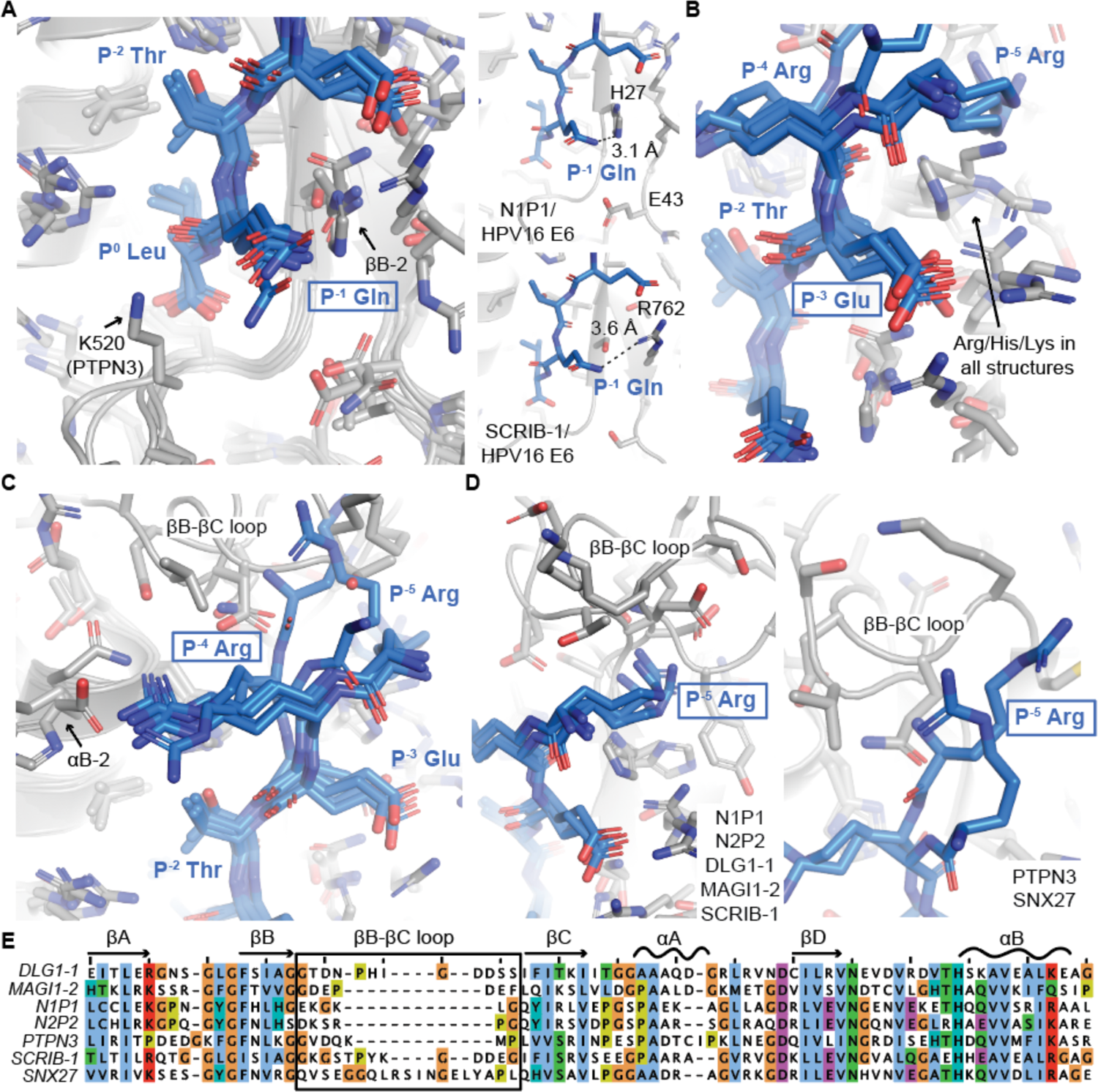
Position-specific interactions with PDZ-peptide models and HPV16 E6. **(A-D**) For all, the PDZ domains are shown in gray cartoon representation, with side chain atoms as sticks and colored by heteroatom (N=blue, O=red). The HPV16 E6 peptides are in blue sticks, colored by heteroatom. Structural features are labeled and described in the main text. (**E**) Sequence alignment of PDZ domains in this analysis was determined using T-coffee multiple sequence alignment, and visualized using Jalview. The variable N- and C-termini are truncated for visual clarity, and secondary structure elements labeled according to MAGI1-2 (6TWQ). The variable βB-βC loop is labeled with a black box.

All seven PDZ domains have a positively-charged residue in the vicinity of the P^-3^ Glu residue in HPV16 E6, suggesting why this is a common preference (**Figure 6B**). Notably, the PTPN3/HPV16 E6 crystal structure shows a stronger interaction between the P^-3^ Glu and hydroxyl of S538 (2.9 Å) as opposed to the amino group of K526 (5.7 Å), which is also nearby.

In all structures and structural models, the P^-4^ Arg is pointed towards the *α*B-2 residue, which is immediately C-terminal to the PDZ Class I conserved *α*B-1 His that interacts with the P^-2^ Thr (**Figure 6C**). For several of these structures, residues in the *β*B-*β*C loop are also located within interaction distances (**Figure 6A**). The P^-5^ Arg interacts directly with *β*B-*β*C loop residues, or is in the vicinity as well (**Figure 6D**). The *β*B-*β*C loop is the most variable region of these PDZ domains, with respect to both sequence and length (**Figure 6E**). This is also the structural element of these PDZ domains that shows the most flexibility (**Figure 6D**), an observation seen in previous PDZ/peptide NMR ensembles (e.g., PDB 4LOB) and elevated temperature factors in most crystallographic structures.

The side-chain atoms of CFTR peptide residues are interacting with similar position-specific amino acids in the PDZ/CFTR structural models. As predicted, N1P1 and N2P2 are the only PDZ domains with a Glu or Asp that directly contacts the P^-1^ Arg (**Figure S2A**). The P^-4^ Gln lies near the *α*B-2 and *β*B-*β*C loop residues (**Figure S2B**). The P^-5^ Val is near the B-*β*C loop, which typically contains polar residues. The exception is SNX27 and V59. The P^-5^ Val is positioned ideally to interact in our model, reflective of the clear P^-5^ Val preference shown in the binding assays, including a 14-fold increase in HPV_P5V_ binding affinity compared to HPV16 E6 as well as 16-fold decrease in binding affinity for CFTR_P5R_ compared to CFTR (**Table 2, Figure S2C**).

The other |*ΔΔ*G°| values that exceed 1 kcal/mol were for SCRIB-1 with HPV_P4Q_ and HPV_P5V_ - HPV16 E6 and PTPN3 with CFTR_P1Q_ - CFTR. In SCRIB-1, the *α*B-2 residue is H794, which likely explains the negative preference for a P^-4^ Arg. In addition, residues in the *β*B-*β*C loop near the P^-5^ position are ^749^TPY^751^, which are relatively hydrophobic and may provide a basis for the 4-fold increase in affinity, 4.5 ± 2.8 μM versus 20 ± 7 μM, upon HPV_P5V_ binding (**Table 2, Figure S2D**).

Electrostatic potential surface maps of all seven PDZ domains show a neutral or slightly negative P^-4^ and P^-5^ binding surface for the PDZ domains with a preference for HPV16 E6, including DLG1-1, MAGI1-2, PTPN3, and SCRIB-1 (**Figure 7**). Conversely, the three PDZ domains with higher relative affinities for CFTR (N1P1, N2P2, and SNX27) are all positively charged in this area and, in general, along the entire binding surface (**Figure 7**). There is a negative surface area on N1P1 and N2P2 due to the Glu/Asp residues that bind the P^-1^ Arg. The PTPN3 surface also shows a positive charge where the P^-5^ position binds, which is consistent with the increased binding affinity for HPV_P5V_, as compared to HPV16 E6 (3.4 μM and 10.1 μM, respectively).

**Figure 7.**
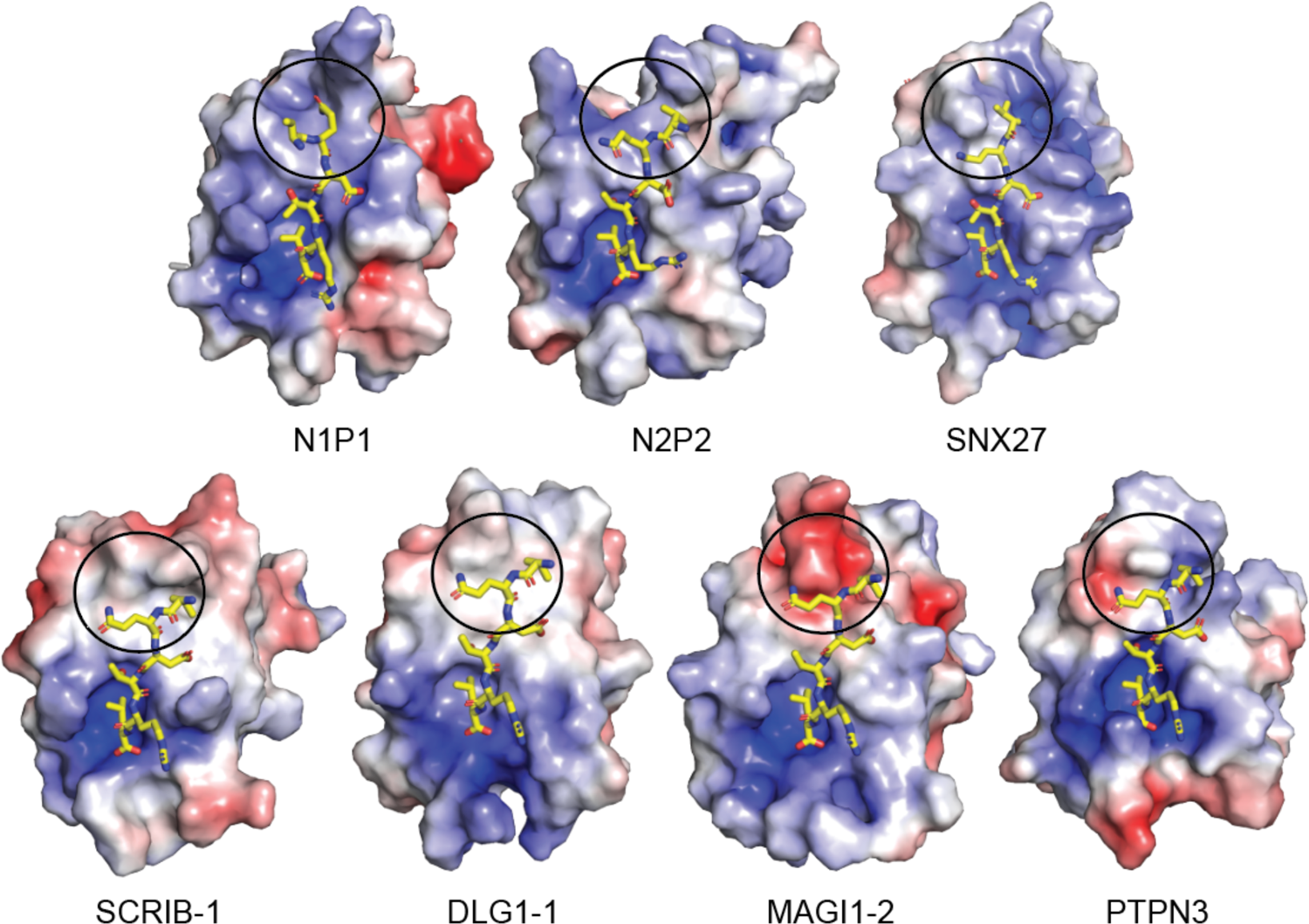
Electrostatic potential surface maps of PDZ/CFTR models. For all, the electrostatic potential surface maps of the PDZ domains from template structures were calculated using APBS in PyMOL. PDZ domains are shown in surface representation, with ± 5 eV represented as a red-to-blue gradient, with red = −5 eV. In general, PDZ domains with a preference for CFTR show a more positive peptide-binding cleft, specifically near the P^-4^ and P^-5^ peptide binding positions, which are both arginine in HPV16 E6. The exception is PTPN3, which does prefer the CFTR P^-5^ Val (**Table 2**), despite an overall preference for HPV16 E6. These areas are highlighted with black circles. PDB IDs used were: N1P1 (1I92), N2P2 (2HE4), SNX27 (6QGL), SCRIB-1 (6MYF), DLG1-1 (3RL7), MAGI1-2 (6TWQ), and PTPN3 (6HKS).

These electrostatic surface potentials in areas of the PDZ domains that interact with upstream peptide residues may also reflect preferences for positions beyond P^-5^. For example, we saw an increased binding affinity for the P^-6^-P^-9^ SSRT residues in _SSRT_CFTR (130 ± 10 μM) as compared to previously determined values for the WT CFTR sequence with P^-6^-P^-9^ TEEE (420 ± 80 μM), consistent with a slightly negatively charged surface (**Figure S3**).^41, 57^ Overall, the experimental structures and our peptide-bound models support the binding data.

### IV. Sequence characteristics of PDZ domains in residues that interact with the P^-1^, P^-4^, and P^-5^ target positions

Finally, we were curious if the positions identified by our structural models as interacting with specific peptide positions are generally applicable to PDZ-peptide complexes. We previously identified seven residues that generally interact with target amino-acid side chains and comprise the peptide-binding cleft.^58^ Here, we wanted to do a similar analysis, but focused on the P^-1^, P^-4^, and P^-5^ positions. We aligned the sequences of 50 Class I PDZ domains in proteins previously identified as HPV16 E6 targets, and which include interacting peptides/proteins. We identified amino acids that interact with each peptide position. The HPV16 E6 and CFTR structural model data described above were largely consistent with this analysis (**Figure S4**).

This alignment revealed that of these domains, the NHERF proteins are the only PDZ domains with the conserved negatively-charged Glu/Asp positioned to interact with the P^-1^ Arg in CFTR. In addition, while the majority of these domains have a polar residue at *α*B-2, several specifically contain a Glu/Asp amino acid, a favorable partner for the P^-4^ Arg in HPV16 E6. Finally, P^-5^ preferences are harder to predict due to the variability in the *β*B-*β*C loops (**Figure S4**). This loop is flexible, including in different molecules of the asymmetric unit, as evidenced by the DLG1-1/APC peptide structure (**Figure S5A**), as well as between PDZ domains. As a selectivity-determining example, the *β*B-*β*C loop of the TIP-1 PDZ domain forms an additional *α*-helix that creates a deep binding pocket, introducing a strong P^-5^ Trp preference (**Figure S5B**). In this set of proteins, one of the shortest *β*B-*β*C loops is in MPDZ-4 at 4 residues, whereas the longest is in MAGI1-5 at 20, highlighting the discrepancy (**Figure S4**). However, in general, an abundance of polar residues at these positions suggests why the HPV16 E6 sequence is broadly well tolerated. These observations suggest that the P^-1^ Arg and P^-5^ Val amino acids in CFTR select against several PDZ targets. In most cases, it is likely a contribution of relatively minor negative preferences at multiple positions in the CFTR sequence that dramatically decrease binding affinities for non-CFTR PDZ targets.

## Concluding Remarks

PDZ domains bind their endogenous targets relatively weakly, with an average binding affinity for calculated values of around 1 μM.^11^ This is biologically relevant, as these interactions are often transient and involved in short-lived trafficking or signaling processes.^9–11^ Another consequence is that the interactome of many of these domains overlaps with other PDZ domains, resulting in a network of both distinct and shared binding preferences and targets.^11^ This characteristic is well exploited by viruses, with several, including SARS-CoV-2, hijacking multiple PDZ domain-mediated processes with a single or just a few pathogenic protein(s).^16, 20, 23–25, 28, 59–61^

Here, we were interested in the hypothesis that viral sequences, e.g., HPV16 E6, may have evolved to be able to target several PDZ proteins, whereas human sequences, e.g., CFTR, bind much fewer due to these varied effects on cellular output. Indeed, we saw that all 28 measured binding affinities for HPV16 E6 and HPV variant peptides are spread relatively evenly from 0.1-100 μM (**Table 2**), a standard range for PDZ-target interactions.^29^ Conversely, the _SSRT_CFTR values revealed a bimodal distribution of relatively high (N1P1, N2P2, SNX27) or low (SCRIB1, DLG1-1, MAGI1-2, PTPN3) affinities, separated by nearly two orders of magnitude. This suggests that CFTR may be under evolutionary selective pressure to be specific for only its biological targets, whereas the viral HPV16 E6 sequence is not. An endogenous sequence that interacts with dozens of PDZ domains may directly and/or indirectly disadvantage critical processes, like receptor trafficking, that are under tight cellular regulation. In contrast, a viral protein that can bind several PDZ domains and integrate quickly into host machinery for, e.g., cell growth and proliferation, may be advantageous. This idea is consistent with the cancer-causing strains of HPV. Notably, there are exceptions to this based on cellular context; for example, CAL PDZ is known to bind its endogenous targets, e.g., CFTR and HPV16 E6, with affinities > 100 μM, suggesting that binding is a result of macromolecular complex formation and increased local concentration, amongst other factors.^14, 34^

Sequences related by evolution support this idea. The C-termini of CFTR sequences related over millions of years of evolution differ only at the P^-3^ and P^-5^ positions, and exhibited only conservative mutations, as underlined, for example, in the sequences for sea lamprey (*Petromyzon marinus*; IQETRL), shark (*Callorhinchus milii*; LQETRL), zebrafish (*Danio rerio*; IQDTRL), chicken (*Gallus gallus*; VQETRL), armadillo (*Dasypus novemcinctus*; VQDTRL), and dog (*Canis lupus familiaris*; VQDTRL), differ only at the P^-3^ and P^-5^ positions, and are limited to conservative mutations, as underlined. In contrast, E6 proteins from different cancer-causing strains of HPV vary more widely, including differences at all positions (underlined), e.g., HPV31 (RTETQV), HPV33 (RRETAL), HPV59 (RSETLV), HPV56 (PRESTV), HPV58 (RRQTQV), and HPV52 (RPVTQV).^28^ This suggests that the CFTR sequence is under higher evolutionary pressure, which we hypothesize is related to its sequence-encoded specificity. Furthermore, endogenous PDZ-peptide interactions often utilize specific additional regulatory mechanisms that further constrain their evolution, e.g., macromolecular complex formation, localization, and/or posttranslational modifications, like phosphorylation.^11^

We specifically chose to study the HPV16 E6 and CFTR sequences because of their striking similarity. Although HPV is known to bind over a dozen PDZ-domain containing proteins, whereas CFTR binds 5-6, the sequences differ mainly in three positions, at P^-1^, P^-4^, and P^-5^, RRETQL and VQDTRL, respectively. Notably, these sequences are identical in the PDZ Class I motif positions, P^0^ and P^-2^. Ultimately, we were surprised to discover that for most HPV16 E6-binding PDZ domains, there is not a single CFTR residue that destroys peptide binding (**Tables 1-2**). The same was true when we tested CFTR variant peptides, most substitutions to HPV16 E6 amino acids at these positions resulted in relatively minor changes in binding affinity (**Table 2**).

There were exceptions with clear structural explanations, specifically positive preferences for a P^-1^ Arg in NHERF PDZ domains, and P^-5^ Val in SNX27 PDZ. The P^-1^ Arg and P^-5^ Val residues are both in CFTR, suggesting that these proteins may have co-evolved to target each other, as previous work suggests.^62^ Overall, our data revealed that the contribution of all three peptide positions appeared to play a role. Indeed, when we summed the WT-compared *ΔΔ*G° values of the three HPV16 E6- or CFTR-substituted peptides for PDZ domains with strong HPV16 E6 (DLG1-1, SCRIB-1, TIP-1, MAGI1-2, PTPN3) or CFTR (N1P1, N2P2) preferences, the |*Σ ΔΔ*G°| value exceeded 1 kcal/mol in all cases, 1.5 kcal/mol for four of the PDZ domains (N2P2, DLG1-1, MAGI1-2, SCRIB-1, TIP-1), and 2 kcal/mol for two of the domains (N1P1, PTPN3) (**Figure 5B**). Our binding data demonstrated that these summed free energies of binding are of the magnitude to explain the 10-60-fold differences in binding affinity measured.

The notion of additive contributions of peptide residues to PDZ binding is well established and based on seminal work in the field by Stiffler *et al.* that characterized the binding selectivity of 157 mouse PDZ domains with respect to 217 genome-encoded preferences.^63^ Several additional groups, using a variety of computational techniques, have also developed methods for predicting the free energy of binding interactions for PDZ domains.^64–69^ Our results used the HPV16 E6 and CFTR PDZ target sequences to experimentally show how this inherent property can also contribute to interaction promiscuity. For the HPV16 E6 viral oncoprotein, a C-terminal sequence that encodes for a large number of PDZ domain targets may be beneficial for propagation of the virus, although it may ultimately be disadvantageous for the host. In contrast, the CFTR C-terminus must be specific for its PDZ targets, such that the protein is properly regulated and trafficked in the cell without off-target effects. Evolutionary pressure towards selectivity for endogenous sequences may also lead to stronger second-order, context-dependent effects in these systems.

It would be interesting to see if viral PDZ-targeting sequences are generally more promiscuous than endogenous partners. For example, there may be evolutionary benefits in the broad targeting of cellular partners. Alternatively, evolution may have targeted a functionally beneficial subset of PDZ domains, but without strong counterselection against off-target sequences. Recent work strongly asserts that climate change will increase the risk of cross-species viral transmission, e.g., SARS-CoV-2, in coming years.^70^ A better understanding of the mechanisms by which viruses infiltrate PDZ domain-mediated pathways may provide therapeutic insight against the oncogenic potential of these infections via direct design of inhibitors for these viral proteins.

## Materials and Methods

### Recombinant protein expression and purification of PDZ domains: CAL, DLG1-1, MAGI1-2, N1P1, N2P2, PTPN3, SCRIB-1, SNX27, and TIP-1

Briefly, all PDZ domains were recombinantly expressed from pET vectors (pET16b for CAL, N1P1, N2P2, and TIP-1, pET28a(+) for all others) in *Escherichia coli* (BL21(DE3) RIL for CAL and TIP-1, BL21(DE3) for all others), and purified using protein chromatography, using identical or similar protocols as those previously described.^34, 36, 41, 51, 71^ Most PDZ domains included His_10_ (CAL, N1P1, N2P2, and TIP-1) or His_6_ (all others) sequences and a HRV 3C (CAL, N1P1, N2P2, and TIP-1) or TEV (all others) protease cleavage site. MAGI1-2 and PTPN3 were additionally expressed as N-terminal SUMO-fusion proteins. PDZ domain boundaries used were: CAL (residues: 284-371), DLG1-1 (220-317), MAGI1-2 (471-554), N1P1 (1-139), N2P2 (142-280), PTPN3 (505-597), SCRIB-1 (722-815), SNX27 (41-135), and TIP-1 (9-120).

Briefly, for DLG1-1, MAGI1-2, PTPN3, and SCRIB-1, which were not described elsewhere by this group, overexpression of the PDZ domain was induced using 0.15 mM IPTG, followed by shaking for 16-20 h at a reduced temperature, 18°C overnight. Centrifuged cells were resuspended in lysis buffer [150 mM NaCl, 50 mM Tris pH 8.5, 10% (*w/v*) glycerol, 10 mM imidazole, 0.5 mM TCEP, Complete EDTA-free protease inhibitor, and DNase], followed by lysis using sonication. Following clarification of the whole cell lysate using centrifugation, the supernatant was collected, filtered using Miracloth (Sigma-Aldrich), and added to a 5 mL NiNTA HisTrap column (Cytiva), equilibrated in 3 column volumes (CV) of wash buffer [150 mM NaCl, 50 mM Tris pH 8.5, 10% (*w/v*) glycerol, 10 mM imidazole, 1 mM TCEP]. The column was washed with 10 CV of wash buffer. Protein was eluted using a gradient from 0-100% of elution buffer [wash buffer with 400 mM imidazole] over 5-10 CV into fractions. Purity was assessed using SDS-PAGE and fractions were concentrated using Amicon 3500 MWCO centrifugal filters (Sigma-Aldrich).

Following concentration, protein was further purified using size exclusion chromatography on a Superdex S75 16/600 (Cytiva) column using SEC buffer [equal to wash buffer without the imidazole]. Purity was determined using SDS-PAGE, relevant fractions collected/concentrated, and final concentrations were determined by measuring absorbance at *λ*=280 nm. Extinction coefficients for N1P1, N2P2, and CAL PDZ were previously experimentally determined, while all others were calculated. Extinction coefficients used were: CAL (2980 M^-1^ cm^-1^), DLG1-1 (4470 M^-1^ cm^-1^), MAGI1-2 (1615 M^-1^ cm^-1^), N1P1 (2980 M^-1^ cm^-1^), N2P2 (2980 M^-1^ cm^-1^), PTPN3 (2980 M^-1^ cm^-1^), SCRIB-1 (8480 M^-1^ cm^-1^), SNX27 (2980 M^-1^ cm^-1^), and TIP-1 (10715 M^-1^ cm^-1^).^34, 41, 51, 58^ Final protein concentrations for CAL were determined by Bradford assay due to CAL PDZ’s tendency to non-specifically bind ATP. Proteins were flash frozen using liquid nitrogen and stored at −80°C.

### Fluorescence polarization assays

Fluorescence polarization assays were performed as previously described, with modifications noted below.^34, 41, 51, 58, 72^ The reporter peptides used were: CAL (*F*\*-iCAL36_Q_, sequence: *FITC*-ANSRWQTSII), DLG1-1 (*F*\*-HPV16 E6, *FITC*-SSRTRRETQL), MAGI1-2 (*F*\*-HPV16 E6), N1P1 (*F\**-CFTR_6_, *FITC*-VQDTRL), N2P2 (*F\**-CFTR_10_, *FITC*-TEEEVQDTRL), PTPN3 (*F*\*-HPV16 E6), SCRIB-1 (*F*\*-HPV16 E6), SNX27 (*F*\*-GIRK3, *FITC*-LPPPESESKV), and TIP-1 (*F*\*-iCAL36). Determined *K*_D_ values were the average calculated from triplicate experiments.^34^ The *K*_D_ values used to determine *K*_I_ competition values were, calculated here (listed ± SD) or previously: CAL (1.03 μM), DLG1-1 (17 ± 4 μM), N1P1 (0.49 μM), N2P2 (0.23 μM), SCRIB-1 (5.1 ± 1.7 μM), SNX27 (0.33 μM), and TIP-1 (0.2 μM). Additional calculated *K*_D_ values in this work were: MAGI1-2 (12 ± 4 μM) and PTPN3 (2 ± 1 μM).

Competition experiments were performed in at least triplicate, with N=11 for SCRIB-1/HPV_P4Q_ due to higher than average variability. Similarly, for DLG1-1, n=3-7 due to experimental variability. The final protein concentrations used for competition experiments were: CAL (1.85 μM), DLG1-1 (15 μM), MAGI1-2 (15 μM), N1P1 (0.75 μM), N2P2 (0.35 μM), PTPN3 (5 μM), SCRIB-1 (7.2 μM), SNX27 (0.5 μM), and TIP-1 (0.36 μM). All assays were conducted using FP buffer [SEC buffer plus 0.1 mg mL^-1^ bovine IgG (Sigma-Aldrich) and 0.5 mM Thesit (Sigma-Aldrich)] Binding affinities were determined using SOLVER, as previously described.^34, 36, 41, 51, 58, 72^ For MAGI1-2 and PTPN3, IC_50_ values were determined using a four-parameter logistic curve fit in Kaleidagraph as previously described.^34^ Kaleidagraph version 5.01 was used to visualize all binding curves.

In contrast to previously published methods, many isotherms were fit using fluorescence polarization values instead of anisotropy, as a technical consequence of the default output of the BioTek Synergy H1 plate reader and version of Gen5 software used. We confirmed that the polarization values track the anisotropy values within a range of ∼4%, and that the effect on computed affinities is less than the standard error of the least-squares fit.

All experiments were conducted at T = 298 K (CAL, TIP-1) or 300 K (all others) and equilibrium free energy values calculated by *Δ*G° = *R*T ln (*K*_I_), where *R*=0.001987 kcal mol^-1^ K^-1^.

### Energy minimization calculations using GROMACS

PDZ-peptide complexes were modeled using PyMOL, using the following PDB codes as templates: 6TWQ (MAGI1-2), 1I92 (N1P1), 2HE4 (N2P2), 6MYF (SCRIB-1), 3RL7 (DLG1-1), 6HKS (PTPN3), and 6QGL (SNX27). Coot was used to build side chains, where missing from the available structure, as well as rotate the orientation of the P^-5^ side chain in SNX27.^73^ PDZ domains were aligned by main chain atoms in order to determine the starting peptide position, using MAGI1-2/HPV16 E6 (6TWQ) as a template. For CFTR peptides, the HPV16 E6 peptide in 6TWQ (or GIRK3 for SNX27/HPV16 E6 and SNX27/CFTR) was mutated using the PyMOL mutagenesis wizard.

The system was solvated using spc216.gro, an equilibrated 3-point solvent model built into GROMACS, as well as a TIP 3-point water model (TIP3P). Ions were added to a 0.15 M concentration, with a neutral net charge (balanced with Na^+^ and Cl^-^), and van der Waals radii were estimated as previously described.^74^ The AMBER99SB-ILDN force field was used and energy minimization conducted using GROMACS version 2022.4.^75–79^ The steepest descent energy minimization was performed on the solvated system with a maximum force tolerance of 1000 kJ mol^-1^ nm^-1^ (239.01 kcal mol^-1^ nm^-1^) for all PDZ-peptide models (**Table S1, Figure S6**).

## Acknowledgements

The authors would like to thank members of the Amacher lab for technical assistance, as well as Dr. Jay McCarty for helpful discussions and assistance with the energy minimizations used for modeling PDZ-peptide complexes, and Dr. Zdenek Svindrych for technical discussions about fluorescence polarization experiments. This work was funded by NSF CHE-1904711 and a Cottrell Scholar Award to JFA. In addition, EFT, MG, and SNR were funded by NSF REU CHE-1757629, and JMB was funded by a Cottrell Postbac Award from the Research Corporation for Science Advancement. NPG and DRM were supported in part by NIH R01-DK101541, P30-DK117469, and P20-GM113132.

## Author contributions

JFA designed the experiments. EFT, JMB, SNJ, MG, NPG, SNS, NJP, SNR, SAS, IGPM, and JFA performed experiments. EFT, JMB, NPG, NJP, DRM, and JFA contributed to data analysis. JFA wrote the manuscript and prepared the figures. All authors contributed to editing of the manuscript.

## Literature Cited

1. Davey NE, Van Roey K, Weatheritt RJ, Toedt G, Uyar B, Altenberg B, Budd A, Diella F, Dinkel H, Gibson TJ (2012) Attributes of short linear motifs. Mol. Biosyst. 8:268–281.

2. Cumberworth A, Lamour G, Babu MM, Gsponer J (2013) Promiscuity as a functional trait: intrinsically disordered regions as central players of interactomes. Biochem. J. 454:361–369.

3. Edwards RJ, Davey NE, O’Brien K, Shields DC (2012) Interactome-wide prediction of short, disordered protein interaction motifs in humans. Mol. Biosyst. 8:282–295.

4. Westermarck J, Ivaska J, Corthals GL (2013) Identification of protein interactions involved in cellular signaling. Mol. Cell Proteomics 12:1752–1763.

5. Kennedy MB (1995) Origin of PDZ (DHR, GLGF) domains. Trends Biochem. Sci. 20:350.

6. Cho KO, Hunt CA, Kennedy MB (1992) The rat brain postsynaptic density fraction contains a homolog of the Drosophila discs-large tumor suppressor protein. Neuron 9:929–942.

7. Woods DF, Bryant PJ (1991) The discs-large tumor suppressor gene of Drosophila encodes a guanylate kinase homolog localized at septate junctions. Cell 66:451–464.

8. Doyle DA, Lee A, Lewis J, Kim E, Sheng M, MacKinnon R (1996) Crystal structures of a complexed and peptide-free membrane protein-binding domain: molecular basis of peptide recognition by PDZ. Cell 85:1067–1076.

9. Nourry C, Grant SGN, Borg J-P (2003) PDZ domain proteins: plug and play! Sci STKE 2003:RE7.

10. Harris BZ, Lim WA (2001) Mechanism and role of PDZ domains in signaling complex assembly. J. Cell Sci. 114:3219–3231.

11. Amacher JF, Brooks L, Hampton TH, Madden DR (2020) Specificity in PDZ-peptide interaction networks: Computational analysis and review. Journal of Structural Biology: X 4:100022.

12. Gujral TS, Karp ES, Chan M, Chang BH, MacBeath G (2013) Family-wide investigation of PDZ domain-mediated protein-protein interactions implicates β-catenin in maintaining the integrity of tight junctions. Chem. Biol. 20:816–827.

13. Kim E, Sheng M (2004) PDZ domain proteins of synapses. Nat. Rev. Neurosci. 5:771–781.

14. Guggino WB, Stanton BA (2006) New insights into cystic fibrosis: molecular switches that regulate CFTR. Nat. Rev. Mol. Cell Biol. 7:426–436.

15. Dunn HA, Ferguson SSG (2015) PDZ Protein Regulation of G Protein-Coupled Receptor Trafficking and Signaling Pathways. Mol. Pharmacol. 88:624–639.

16. Javier RT, Rice AP (2011) Emerging theme: cellular PDZ proteins as common targets of pathogenic viruses. J. Virol. 85:11544–11556.

17. Shepley-McTaggart A, Sagum CA, Oliva I, Rybakovsky E, DiGuilio K, Liang J, Bedford MT, Cassel J, Sudol M, Mullin JM, et al. (2021) SARS-CoV-2 Envelope (E) protein interacts with PDZ-domain-2 of host tight junction protein ZO1. PLoS One 16:e0251955.

18. Saidu NEB, Filić V, Thomas M, Sarabia-Vega V, Đukić A, Miljković F, Banks L, Tomaić V (2019) PDZ Domain-Containing Protein NHERF-2 Is a Novel Target of Human Papillomavirus 16 (HPV-16) and HPV-18. J. Virol. 94.

19. Caillet-Saguy C, Durbesson F, Rezelj VV, Gogl G, Dinh Tran Q, Twizere J-C, Vignuzzi M, Vincentelli R, Wolff N (2021) Host PDZ-containing proteins targeted by SARS-Cov-2. FEBS J.

20. James CD, Roberts S (2016) Viral Interactions with PDZ Domain-Containing Proteins-An Oncogenic Trait? Pathogens 5.

21. White EA, Kramer RE, Tan MJA, Hayes SD, Harper JW, Howley PM (2012) Comprehensive analysis of host cellular interactions with human papillomavirus E6 proteins identifies new E6 binding partners and reflects viral diversity. J. Virol. 86:13174–13186.

22. Liu Y, Henry GD, Hegde RS, Baleja JD (2007) Solution structure of the hDlg/SAP97 PDZ2 domain and its mechanism of interaction with HPV-18 papillomavirus E6 protein. Biochemistry 46:10864–10874.

23. Pim D, Bergant M, Boon SS, Ganti K, Kranjec C, Massimi P, Subbaiah VK, Thomas M, Tomaić V, Banks L (2012) Human papillomaviruses and the specificity of PDZ domain targeting. FEBS J. 279:3530–3537.

24. Zhang Q, Gefter J, Sneddon WB, Mamonova T, Friedman PA (2021) ACE2 interaction with cytoplasmic PDZ protein enhances SARS-CoV-2 invasion. iScience 24:102770.

25. Ivarsson Y, Arnold R, McLaughlin M, Nim S, Joshi R, Ray D, Liu B, Teyra J, Pawson T, Moffat J, et al. (2014) Large-scale interaction profiling of PDZ domains through proteomic peptide-phage display using human and viral phage peptidomes. Proc. Natl. Acad. Sci. USA 111:2542–2547.

26. Thomas M, Kranjec C, Nagasaka K, Matlashewski G, Banks L (2011) Analysis of the PDZ binding specificities of Influenza A virus NS1 proteins. Virol. J. 8:25.

27. Thomas M, Myers MP, Massimi P, Guarnaccia C, Banks L (2016) Analysis of Multiple HPV E6 PDZ Interactions Defines Type-Specific PDZ Fingerprints That Predict Oncogenic Potential. PLoS Pathog. 12:e1005766.

28. Ganti K, Broniarczyk J, Manoubi W, Massimi P, Mittal S, Pim D, Szalmas A, Thatte J, Thomas M, Tomaić V, et al. (2015) The human papillomavirus E6 PDZ binding motif: from life cycle to malignancy. Viruses 7:3530–3551.

29. Vincentelli R, Luck K, Poirson J, Polanowska J, Abdat J, Blémont M, Turchetto J, Iv F, Ricquier K, Straub M-L, et al. (2015) Quantifying domain-ligand affinities and specificities by high-throughput holdup assay. Nat. Methods 12:787–793.

30. Duhoo Y, Girault V, Turchetto J, Ramond L, Durbesson F, Fourquet P, Nominé Y, Cardoso V, Sequeira AF, Brás JLA, et al. (2019) High-Throughput Production of a New Library of Human Single and Tandem PDZ Domains Allows Quantitative PDZ-Peptide Interaction Screening Through High-Throughput Holdup Assay. Methods Mol. Biol. 2025:439–476.

31. Ren A, Zhang W, Yarlagadda S, Sinha C, Arora K, Moon C-S, Naren AP (2013) MAST205 competes with cystic fibrosis transmembrane conductance regulator (CFTR)-associated ligand for binding to CFTR to regulate CFTR-mediated fluid transport. J. Biol. Chem. 288:12325–12334.

32. McDermott MI, Thelin WR, Chen Y, Lyons PT, Reilly G, Gentzsch M, Lei C, Hong W, Stutts MJ, Playford MP, et al. (2018) Sorting nexin 27 (SNX27): A novel regulator of cystic fibrosis transmembrane conductance regulator (CFTR) trafficking. BioRxiv.

33. Swiatecka-Urban A, Duhaime M, Coutermarsh B, Karlson KH, Collawn J, Milewski M, Cutting GR, Guggino WB, Langford G, Stanton BA (2002) PDZ domain interaction controls the endocytic recycling of the cystic fibrosis transmembrane conductance regulator. J. Biol. Chem. 277:40099–40105.

34. Cushing PR, Fellows A, Villone D, Boisguérin P, Madden DR (2008) The relative binding affinities of PDZ partners for CFTR: a biochemical basis for efficient endocytic recycling. Biochemistry 47:10084–10098.

35. Cheng J, Moyer BD, Milewski M, Loffing J, Ikeda M, Mickle JE, Cutting GR, Li M, Stanton BA, Guggino WB (2002) A Golgi-associated PDZ domain protein modulates cystic fibrosis transmembrane regulator plasma membrane expression. J. Biol. Chem. 277:3520–3529.

36. Amacher JF, Cushing PR, Brooks L, Boisguerin P, Madden DR (2014) Stereochemical preferences modulate affinity and selectivity among five PDZ domains that bind CFTR: comparative structural and sequence analyses. Structure 22:82–93.

37. Tonikian R, Zhang Y, Sazinsky SL, Currell B, Yeh J-H, Reva B, Held HA, Appleton BA, Evangelista M, Wu Y, et al. (2008) A specificity map for the PDZ domain family. PLoS Biol. 6:e239.

38. Jeong KW, Kim HZ, Kim S, Kim YS, Choe J (2007) Human papillomavirus type 16 E6 protein interacts with cystic fibrosis transmembrane regulator-associated ligand and promotes E6-associated protein-mediated ubiquitination and proteasomal degradation. Oncogene 26:487–499.

39. Hampson L, Li C, Oliver AW, Kitchener HC, Hampson IN (2004) The PDZ protein Tip-1 is a gain of function target of the HPV16 E6 oncoprotein. Int. J. Oncol. 25:1249–1256.

40. Drews CM, Case S, Vande Pol SB (2019) E6 proteins from high-risk HPV, low-risk HPV, and animal papillomaviruses activate the Wnt/β-catenin pathway through E6AP-dependent degradation of NHERF1. PLoS Pathog. 15:e1007575.

41. Amacher JF, Cushing PR, Bahl CD, Beck T, Madden DR (2013) Stereochemical determinants of C-terminal specificity in PDZ peptide-binding domains: a novel contribution of the carboxylate-binding loop. J. Biol. Chem. 288:5114–5126.

42. Karthikeyan S, Leung T, Ladias JA (2001) Structural basis of the Na+/H+ exchanger regulatory factor PDZ1 interaction with the carboxyl-terminal region of the cystic fibrosis transmembrane conductance regulator. J. Biol. Chem. 276:19683–19686.

43. Vistrup-Parry M, Sneddon WB, Bach S, Strømgaard K, Friedman PA, Mamonova T (2021) Multisite NHERF1 phosphorylation controls GRK6A regulation of hormone-sensitive phosphate transport. J. Biol. Chem. 296:100473.

44. Kiyono T, Hiraiwa A, Fujita M, Hayashi Y, Akiyama T, Ishibashi M (1997) Binding of high-risk human papillomavirus E6 oncoproteins to the human homologue of the Drosophila discs large tumor suppressor protein. Proc. Natl. Acad. Sci. USA 94:11612–11616.

45. Yoshimatsu Y, Nakahara T, Tanaka K, Inagawa Y, Narisawa-Saito M, Yugawa T, Ohno S-I, Fujita M, Nakagama H, Kiyono T (2017) Roles of the PDZ-binding motif of HPV 16 E6 protein in oncogenic transformation of human cervical keratinocytes. Cancer Sci. 108:1303–1309.

46. Genera M, Samson D, Raynal B, Haouz A, Baron B, Simenel C, Guerois R, Wolff N, Caillet-Saguy C (2019) Structural and functional characterization of the PDZ domain of the human phosphatase PTPN3 and its interaction with the human papillomavirus E6 oncoprotein. Sci. Rep. 9:7438.

47. Nguyen ML, Nguyen MM, Lee D, Griep AE, Lambert PF (2003) The PDZ ligand domain of the human papillomavirus type 16 E6 protein is required for E6’s induction of epithelial hyperplasia in vivo. J. Virol. 77:6957–6964.

48. Ganti K, Massimi P, Manzo-Merino J, Tomaić V, Pim D, Playford MP, Lizano M, Roberts S, Kranjec C, Doorbar J, et al. (2016) Interaction of the Human Papillomavirus E6 Oncoprotein with Sorting Nexin 27 Modulates Endocytic Cargo Transport Pathways. PLoS Pathog. 12:e1005854.

49. Webb Strickland S, Brimer N, Lyons C, Vande Pol SB (2018) Human Papillomavirus E6 interaction with cellular PDZ domain proteins modulates YAP nuclear localization. Virology 516:127–138.

50. Gogl G, Jane P, Caillet-Saguy C, Kostmann C, Bich G, Cousido-Siah A, Nyitray L, Vincentelli R, Wolff N, Nomine Y, et al. (2020) Dual Specificity PDZ- and 14-3-3-Binding Motifs: A Structural and Interactomics Study. Structure 28:747–759.e3.

51. Gao M, Mackley IGP, Mesbahi-Vasey S, Bamonte HA, Struyvenberg SA, Landolt L, Pederson NJ, Williams LI, Bahl CD, Brooks L, et al. (2020) Structural characterization and computational analysis of PDZ domains in Monosiga brevicollis. Protein Sci. 29:2226–2244.

52. Lauffer BEL, Melero C, Temkin P, Lei C, Hong W, Kortemme T, von Zastrow M (2010) SNX27 mediates PDZ-directed sorting from endosomes to the plasma membrane. J. Cell Biol. 190:565–574.

53. Balana B, Maslennikov I, Kwiatkowski W, Stern KM, Bahima L, Choe S, Slesinger PA (2011) Mechanism underlying selective regulation of G protein-gated inwardly rectifying potassium channels by the psychostimulant-sensitive sorting nexin 27. Proc. Natl. Acad. Sci. USA 108:5831– 5836.

54. Elkins JM, Papagrigoriou E, Berridge G, Yang X, Phillips C, Gileadi C, Savitsky P, Doyle DA (2007) Structure of PICK1 and other PDZ domains obtained with the help of self-binding C-terminal extensions. Protein Sci. 16:683–694.

55. Awadia S, Huq F, Arnold TR, Goicoechea SM, Sun YJ, Hou T, Kreider-Letterman G, Massimi P, Banks L, Fuentes EJ, et al. (2019) SGEF forms a complex with Scribble and Dlg1 and regulates epithelial junctions and contractility. J. Cell Biol. 218:2699–2725.

56. Zhang Z, Li H, Chen L, Lu X, Zhang J, Xu P, Lin K, Wu G (2011) Molecular basis for the recognition of adenomatous polyposis coli by the Discs Large 1 protein. PLoS One 6:e23507.

57. Amacher JF, Zhao R, Spaller MR, Madden DR (2014) Chemically modified peptide scaffolds target the CFTR-associated ligand PDZ domain. PLoS One 9:e103650.

58. Wofford HA, Myers-Dean J, Vogel BA, Alamo KAE, Longshore-Neate FA, Jagodzinski F, Amacher JF (2021) Domain Analysis and Motif Matcher (DAMM): A Program to Predict Selectivity Determinants in Monosiga brevicollis PDZ Domains Using Human PDZ Data. Molecules 26.

59. Kim JI, Hwang M-W, Lee I, Park S, Lee S, Bae J-Y, Heo J, Kim D, Jang S-I, Park MS, et al. (2014) The PDZ-binding motif of the avian NS1 protein affects transmission of the 2009 influenza A(H1N1) virus. Biochem. Biophys. Res. Commun. 449:19–25.

60. Rice AP, Kimata JT (2021) SARS-CoV-2 likely targets cellular PDZ proteins: a common tactic of pathogenic viruses. Future Virol.

61. Gutiérrez-González LH, Santos-Mendoza T (2019) Viral targeting of PDZ polarity proteins in the immune system as a potential evasion mechanism. FASEB J. 33:10607–10617.

62. Ernst A, Gfeller D, Kan Z, Seshagiri S, Kim PM, Bader GD, Sidhu SS (2010) Coevolution of PDZ domain-ligand interactions analyzed by high-throughput phage display and deep sequencing. Mol. Biosyst. 6:1782–1790.

63. Stiffler MA, Chen JR, Grantcharova VP, Lei Y, Fuchs D, Allen JE, Zaslavskaia LA, MacBeath G (2007) PDZ domain binding selectivity is optimized across the mouse proteome. Science 317:364–369.

64. Kamisetty H, Ghosh B, Langmead CJ, Bailey-Kellogg C (2014) Learning Sequence Determinants of Protein:protein Interaction Specificity with Sparse Graphical Models. Res. Comput. Mol. Biol. 8394:129–143.

65. Thomas J, Ramakrishnan N, Bailey-Kellogg C (2009) Graphical models of protein-protein interaction specificity from correlated mutations and interaction data. Proteins 76:911–929.

66. Shao X, Tan CSH, Voss C, Li SSC, Deng N, Bader GD (2011) A regression framework incorporating quantitative and negative interaction data improves quantitative prediction of PDZ domain-peptide interaction from primary sequence. Bioinformatics 27:383–390.

67. Kaufmann K, Shen N, Mizoue L, Meiler J (2011) A physical model for PDZ-domain/peptide interactions. J Mol Model 17:315–324.

68. Holt GT, Jou JD, Gill NP, Lowegard AU, Martin JW, Madden DR, Donald BR (2019) Computational Analysis of Energy Landscapes Reveals Dynamic Features That Contribute to Binding of Inhibitors to CFTR-Associated Ligand. J. Phys. Chem. B 123:10441–10455.

69. Roberts KE, Cushing PR, Boisguerin P, Madden DR, Donald BR (2012) Computational design of a PDZ domain peptide inhibitor that rescues CFTR activity. PLoS Comput. Biol. 8:e1002477.

70. Carlson CJ, Albery GF, Merow C, Trisos CH, Zipfel CM, Eskew EA, Olival KJ, Ross N, Bansal S (2022) Climate change increases cross-species viral transmission risk. Nature 607:555– 562.

71. Valgardson JD, Struyvenberg SA, Sailer ZR, Piper IM, Svendsen JE, Johnson DA, Vogel BA, Antos JM, Harms MJ, Amacher JF (2022) Comparative Analysis and Ancestral Sequence Reconstruction of Bacterial Sortase Family Proteins Generates Functional Ancestral Mutants with Different Sequence Specificities. Bacteria 1:121–135.

72. Valgardson J, Cosbey R, Houser P, Rupp M, Van Bronkhorst R, Lee M, Jagodzinski F, Amacher JF (2019) MotifAnalyzer-PDZ: A computational program to investigate the evolution of PDZ-binding target specificity. Protein Sci. 28:2127–2143.

73. Emsley P, Lohkamp B, Scott WG, Cowtan K (2010) Features and development of Coot. Acta Crystallogr. Sect. D, Biol. Crystallogr. 66:486–501.

74. Bondi A (1964) van der Waals Volumes and Radii. J. Phys. Chem. 68:441–451.

75. Lindorff-Larsen K, Piana S, Palmo K, Maragakis P, Klepeis JL, Dror RO, Shaw DE (2010) Improved side-chain torsion potentials for the Amber ff99SB protein force field. Proteins 78:1950–1958.

76. Van Der Spoel D, Lindahl E, Hess B, Groenhof G, Mark AE, Berendsen HJC (2005) GROMACS: fast, flexible, and free. J. Comput. Chem. 26:1701–1718.

77. Berendsen HJC, van der Spoel D, van Drunen R (1995) GROMACS: A message-passing parallel molecular dynamics implementation. Comput Phys Commun 91:43–56.

78. Wang J, Wolf RM, Caldwell JW, Kollman PA, Case DA (2004) Development and testing of a general amber force field. J. Comput. Chem. 25:1157–1174.

79. Abraham MJ, Murtola T, Schulz R, Páll S, Smith JC, Hess B, Lindahl E (2015) GROMACS: High performance molecular simulations through multi-level parallelism from laptops to supercomputers. SoftwareX 1–2:19–25.

