## Supplemental Material for "Additive energetic contributions of multiple peptide positions determine the relative promiscuity of viral and human sequences for PDZ domain targets"

by

Elise F. Tahti, Jadon M. Blount, Sophie N. Jackson, Melody Gao, Nicholas P. Gill, Sarah N. Smith, Nick J. Pederson, Simone N. Rumph, Sarah A. Struyvenberg, Iain G. P. Mackley, Dean R. Madden, Jeanine F. Amacher

**Table of contents:**

**Figure S1**. PDZ domain template structures for peptide-bound models. 2

**Figure S2.** Position-specific interactions in CFTR-bound models. 3

**Figure S3.** Electrostatic potential surface map of CAL PDZ. 4

**Figure S4.** Sequence alignment of 53 PDZ domains with peptide-bound structures in the Protein Data Bank. 5

**Figure S5.** The βB-βC varies widely in confirmation between PDZ domains and contributes directly to selectivity. 6

**Table S1.** Energy minimization data. 7

**Figure S6.** Energy minimization graphs. 8

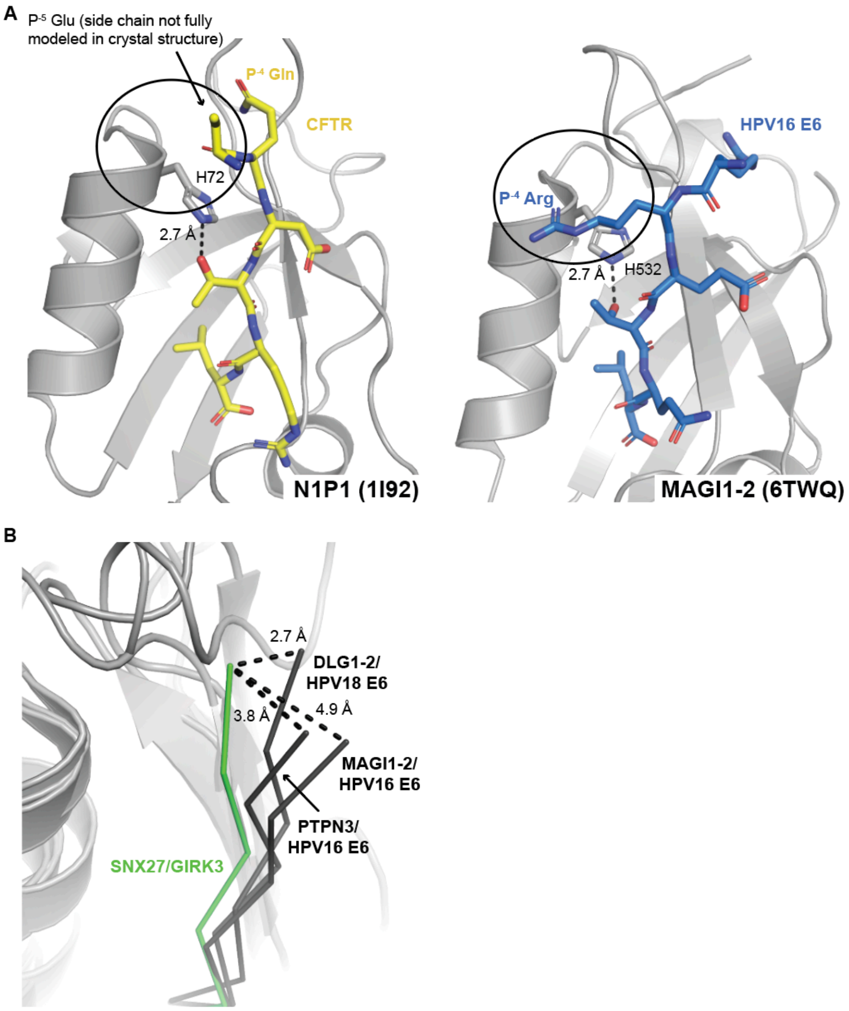


**Figure S1. PDZ domain template structures for peptide-bound models.** All structures are rendered with the PDZ domain shown as gray cartoon. Relevant side chain atoms are shown as sticks, colored by heteroatom (N=blue, O=red), and labeled. Specifically, the conserved αB-1 His, which forms a hydrogen bond with the $P^{-2}$ Ser/Thr for Class I PDZ domains is shown. (**A**) Likely due to the nature of the crystallization conditions, wherein the CFTR peptide (yellow sticks, colored by heteroatom) is a C-terminal extension and binding an N1P1 molecule related by symmetry, the $P^{-4}$ Gln is interacting in the characteristic $P^{-5}$ pocket, and the $P^{-5}$ Glu is interacting with solvent (left). This region is highlighted with a black circle. For comparison, the MAGI1-2/HPV16 E6 structure is shown (right) with the peptide as blue sticks, colored by heteroatom. The same region is circled. (**B**) The confirmation of the βB-βC loop in SNX27 (PDB ID 6QGL) results in the $C_\alpha$ atom of the $P^{-5}$ position is shifted over 2 Å from other peptide-bound PDZ structures. Here, the peptides are shown as ribbon traces, with GIRK3 colored green, and other peptides colored black. DLG1-2/HPV18 E6 (2OQS) is used as an additional comparison.

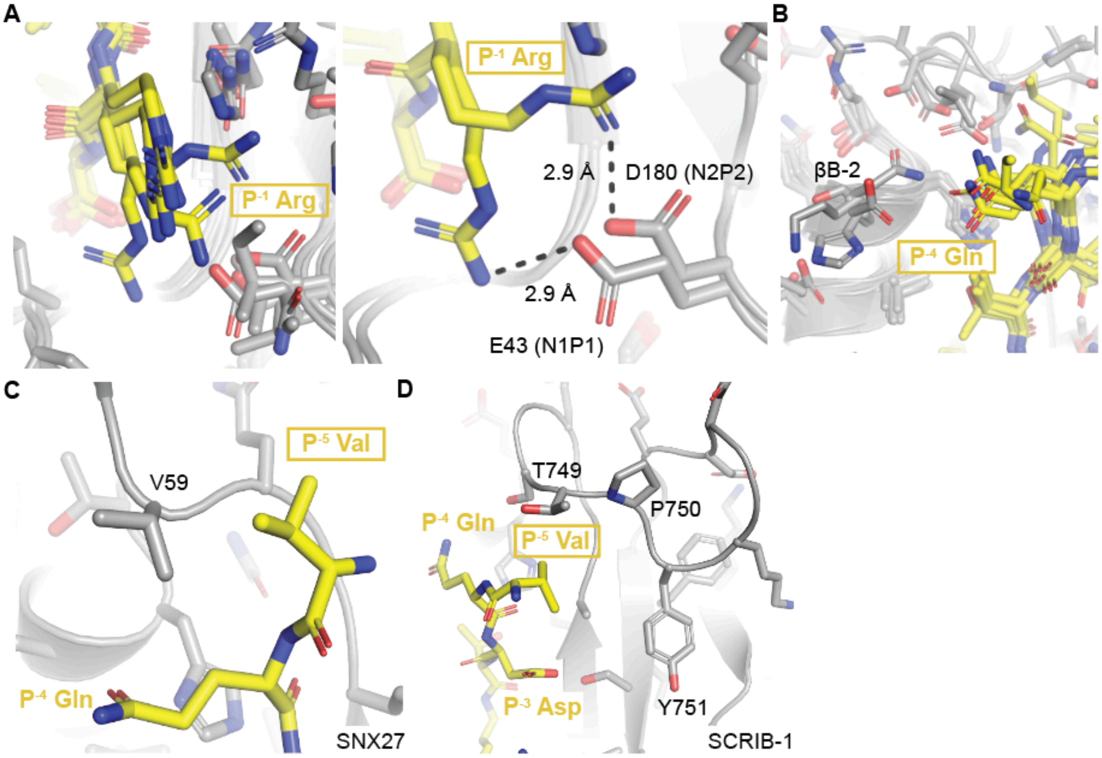


**Figure S2. Position-specific interactions in CFTR-bound models.** All structures are rendered equivalently here, with the PDZ domain shown in cartoon representation, with side chain atoms colored by heteroatom (N=blue, O=red). The CFTR peptides are shown as yellow sticks, colored by heteroatom. In (**A**, right), the distances of electrostatic interactions with E43 (N1P1) and D180 (N2P2) are labeled.

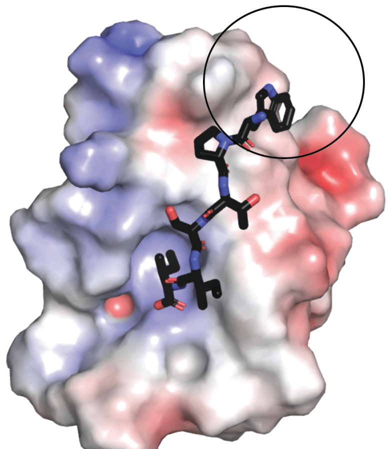


**Figure S3. Electrostatic potential surface map of CAL PDZ.** The electrostatic potential surface of CAL PDZ domain (PDB ID 4E34) is shown bound to the inhibitor iCAL36 peptide, in black sticks and colored by heteroatom (N=blue, O=red). The sequence of iCAL36 is WPTSII, with the $P^{-5}$ Trp circled. The surface of CAL that likely interacts with target residues at $P^{-5}$ or upstream is slightly negatively charged, reflective of the increased binding affinity in $_{SSRT}$CFTR ($P^{-6}$-$P^{-9}$ = SSRT) compared to WT CFTR ($P^{-6}$-$P^{-9}$ = TEEE). The charge here is ± 5 eV represented as a red-to-blue gradient, with red = -5 eV.

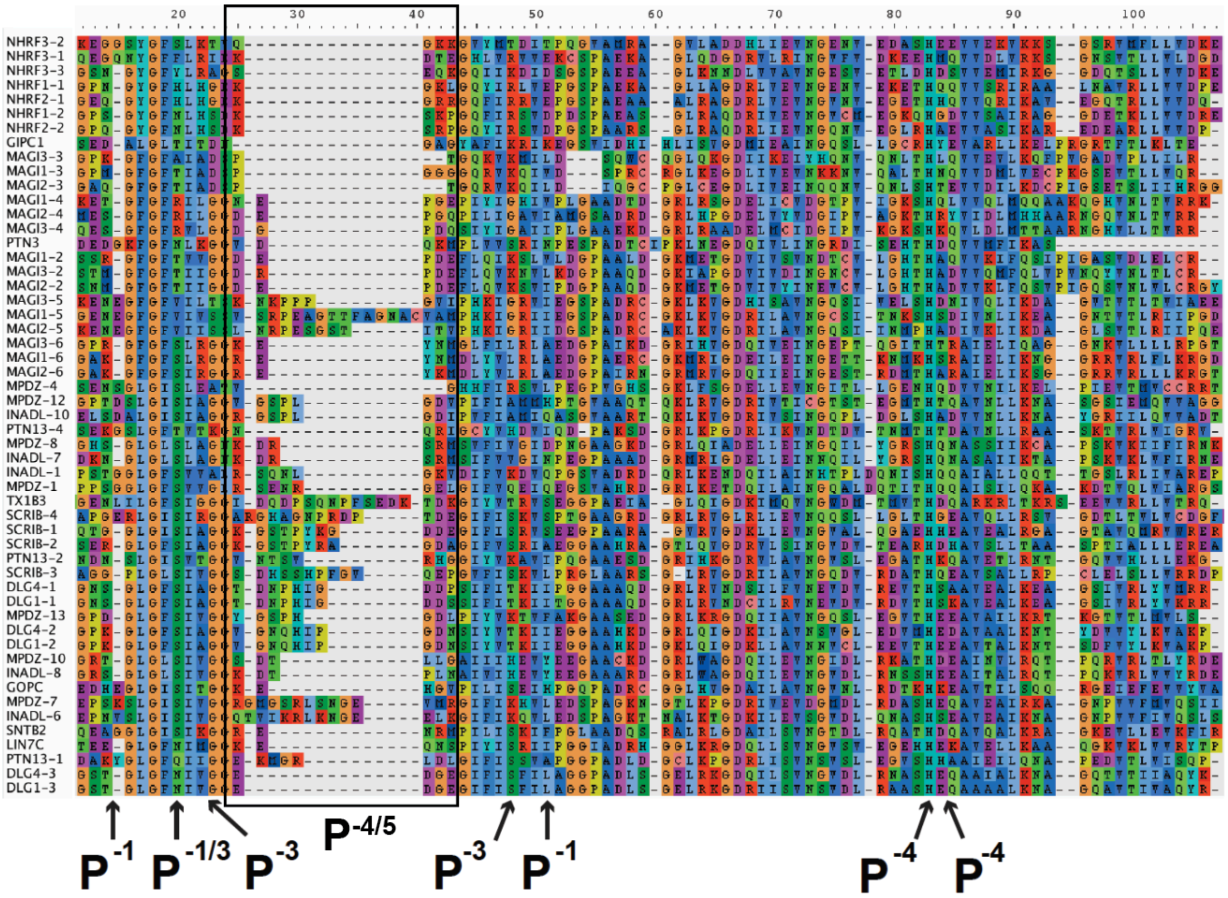


**Figure S4. Sequence alignment of 53 PDZ domains with peptide-bound structures in the Protein Data Bank (PDB).** Sequence alignment of 53 PDZ domain sequences that have peptide-bound structures in the PDB. All structures were analyzed for peptide-binding residues, as labeled above. Sequence alignment was performed using T-coffee, and visualization done in Jalview.

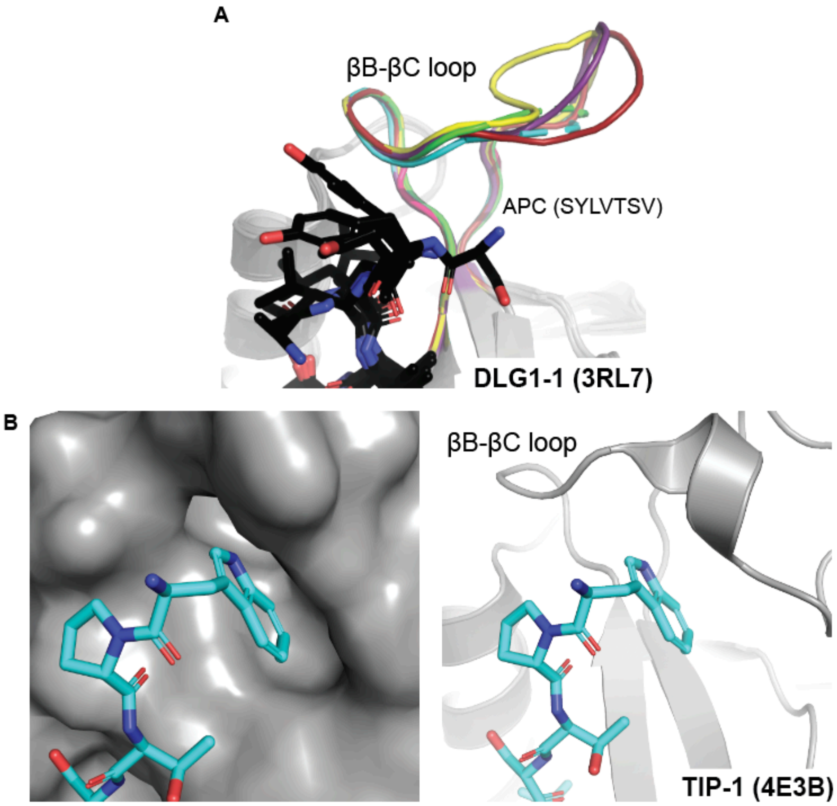


**Figure S5. The βB-βC varies widely in confirmation between PDZ domains and contributes directly to selectivity.** (**A**) Different molecules in the asymmetric unit of DLG1-1/APC (PDB ID 3RL7) reveal differences in the βB-βC loop confirmation, including molecules where the full loop is not resolved. DLG1-1 is shown as a gray cartoon, with the βB-βC loops highlighted using different colors. The APC peptide is in black sticks, colored by heteroatom (N=blue, O=red). (**B**) The βB-βC loop in TIP-1 forms an α-helix that creates a Trp/aromatic-specific pocket. Here, TIP-1 is bound to $iCAL36_L$, an engineered peptide to inhibit CAL. On the left, TIP-1 is in gray surface. On the right, TIP-1 is shown as gray cartoon. The peptide is in cyan sticks, colored by heteroatom.

**Table S1. Energy minimization data.**

| Name[a] | Initial Energy (kJ/mol) | After Energy Minimization (kJ/mol) | Change in Energy (kJ/mol) | Number of Steps |
|---|---|---|---|---|
| DLG1-1_CFTR | -121573.5703 | -353070.7188 | -231497.1485 | 681 |
| MAGI1-2_CFTR | -158210.4844 | -385796.4688 | -227585.9844 | 600 |
| N1P1_HPV | 7399443456 | -360066.6875 | -7399803523 | 518 |
| N2P2_CFTR | -157950 | -473788.9375 | -315838.9375 | 677 |
| N2P2_HPV | 19963038 | -455865.1563 | -20418903.16 | 546 |
| PTPN3_CFTR | -122957.1563 | -365749.9063 | -242792.75 | 594 |
| SCRIB-1_CFTR | -123616.3359 | -348963.6563 | -225347.3204 | 705 |
| SCRIB-1_HPV | 15009923 | -341682.5313 | -15351605.53 | 608 |
| SNX27_CFTR | -113825.9375 | -446543.1875 | -332717.25 | 763 |
| SNX27_HPV | -140083.6719 | -449494 | -309410.3281 | 844 |

[a]PDB IDs used:
DLG1-1 (PDB ID 3RL7)
MAGI1-2 (6TWQ)
N1P1 (1I92)
N2P2 (2HE4)
PTPN3 (6HKS)
SCRIB-1 (6MYF)
SNX27 (6QGL)

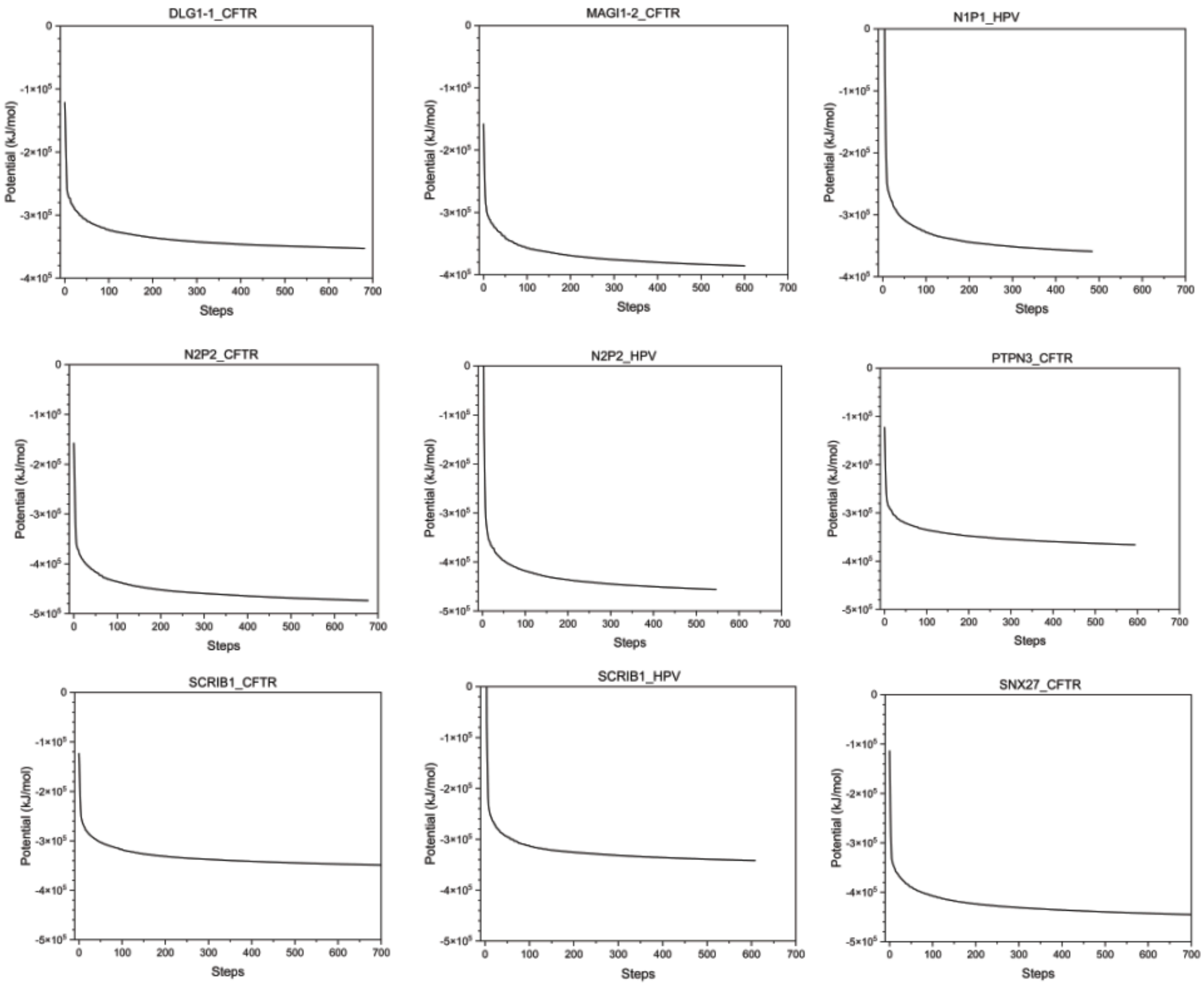


**Figure S6. Energy minimization graphs.** Energy minimization of the PDZ-peptide structures solvated in water. These graphs indicate how many steps were required for each to achieve energy minimization of the complexes.
